# Robust retrieval of data stored in DNA by de Bruijn graph-based *de novo* strand assembly

**DOI:** 10.1101/2020.12.20.423642

**Authors:** Lifu Song, Feng Geng, Ziyi Gong, Xin Chen, Jijun Tang, Chunye Gong, Libang Zhou, Rui Xia, Mingzhe Han, Jingyi Xu, Bingzhi Li, Yingjin Yuan

## Abstract

DNA data storage is a rapidly developing technology with great potential due to its high density, long-term durability, and low maintenance cost. The major technical challenges include various errors, such as the strand breaks, rearrangements, and indels that frequently arise during DNA synthesis, amplification, sequencing, and preservation. Through a *de novo* assembly strategy, we developed an algorithm based on the de Bruijn graph and greedy path search (DBGPS) to address these issues. DBGPS shows distinct advantages in handling DNA breaks, rearrangements, and indels. The robustness of DBGPS is demonstrated by accelerated aging, multiple independent data retrievals, deep error-prone PCR, and large data scale simulations. Remarkably, 6.8 MB of data can be retrieved accurately from a seriously corrupted sample that has been treated at 70 °C for 70 days. With DBGPS, we were able to achieve a logical density of 1.30 bits/cycle and a physical density of 295 PB/g.

**One-Sentence Summary:** A de Bruijn graph-based *de novo* assembly algorithm for DNA data storage enables fast and robust data readouts even with DNA samples that have been severely corrupted.

## Introduction

DNA is the natural solution for the preservation of the genetic information of all life forms on earth. Recently, million years old genomic DNA of mammoths was successfully decoded, revealing its great potential as a long-term data carrier under frozen conditions^1^. Owing to its high density, and low maintenance cost, revealed by recent studies, DNA has been considered as an ideal storage medium to meet the emerging challenge of data explosion^2–20^. Frequently occurring errors in DNA synthesis, amplification, sequencing, and preservation, however, challenge the data reliability in DNA. Efforts to solve these issues led to the implementation of a codec system that requires two layers of error correction (EC) codes, the outer layer codes handling the strand dropouts and the inner layer codes dealing with intramolecular errors, ensuring accurate data readouts^21–25^. The design of outer codes is straightforward since strand dropouts can be well solved by sophisticated erasure codes, *e*.*g*. Fountain, Reed-Solomon (RS) codes^21–23^. In contrast, the design of an inner codec system is challenging due to the complex errors and the unique feature of ‘data reputations’, *i*.*e*., the noisy strand copies^10,21,26^. The traditional decoding process of inner codes generally consists of two steps: clustering and strand reconstruction^23,26,27^. Strand reconstruction from its error-rich copies is a trace reconstruction problem introduced two decades ago^28^. Although there are many theoretical studies on the trace reconstruction problem, practical trace reconstruction algorithms are surprisingly lacking, especially for DNA data storage^26,29–33^. Most studies on trace reconstruction are motivated by the multiple-alignment problem in computational biology^23,28–31,34^. The trace reconstruction problem with the error types of substitutions and indels has been studied, revealing the difficulties of handling indels^23,28–31,34^. In practice, however, DNA breaks and rearrangements occur frequently during the preservation of DNA molecules and PCR-based data copying, threatening the robustness of DNA data storage.

DNA is subjected to hydrolysis and degradation under certain conditions and in long-term storage, causing DNA breaks. This highlights the importance of data stability studies on the influences of harsh conditions, such as high temperature or UV exposure, and methods of DNA protection^21,35,36^. DNA rearrangements refer to the breakage and rejoining of DNA strands. Unspecific amplification, a typical problem with PCR, is the main source of DNA rearrangements in DNA data storage. People currently carefully design primers and optimize PCR conditions to avoid unspecific amplification and employ gel purification to physically exclude rearranged strands from the data pool^23^. An inner codec system that can tolerate fragmented and rearranged DNA strands is nevertheless critical for enhancing the robustness of DNA data storage. The clustering^27,37^ and multiple-alignment^38–40^ algorithms are both incapable of dealing with DNA breaks and rearrangements. A new mechanism that can efficiently handle DNA breaks and rearrangements is highly desirable.

In this study, we proposed a *de novo* assembly-based strategy to handle the complex errors, especially the DNA breaks and rearrangements, in DNA data storage channel. Different from the clustering and multiple alignment (CL-MA) based methods, we first decompose all the strand sequences into *k*-mers with de Bruijn graph (DBG) theory^41–44^. Then, low-occurrence *k*-mers were excluded, omitting huge errors. After that, the strands are assembled straightforwardly by greedy path search and path selection with the aid of the embedded redundancy codes. Compared to the CL-MA-based methods, this DBG based greedy path search algorithm (DBGPS), shows substantial advantages in the handling of errors, especially DNA breaks, rearrangements, and indels. We also verified the effectiveness of DBGPS by simulations up to 1 GB (Giga Bytes, 10^9^ Bytes) scale. The robustness of DBGPS was further proved by three harsh experiments of deep error-prone PCR, multiple data retrievals, and accelerated aging experiments at a data scale of 6.8 MB (Mega Bytes, 10^6^ Bytes). Remarkably, we are able to precisely retrieve the entire 6.8 MB data from a DNA solution that has been incubated at 70 °C for 70 days without any particular protection using DBGPS. Besides the high data robustness, we were able to achieve a high logical density of 1.3 bits/synthesis cycle and a high physical density of 295 PB/g (Peta Bytes, 10^15^ Bytes) using DBGPS.

## Results

### Design of DBGPS: a *de novo* assembly-based strand reconstruction algorithm

The DNA data storage channel is a complex channel with types of errors. Most previous studies have focused on substitution and indel errors^28,30,36,45–48^. In practice, however, DNA breaks and rearrangements occur frequently during the preservation of DNA molecules and PCR-based data copying, threatening the robustness of DNA data storage. These two types of errors should be considered in the DNA data storage channel (**Fig. S1A**). However, DNA breaks and rearrangements can cause severe failures to the clustering and multiple-alignment algorithms. To handle these issues, as illustrated in **Fig. 1A**, we focus on a distinct route of *de novo* assembly which can potentially take advantage of the “multi-copy” feature of DNAs for accurate strand reconstruction. The successful applications of de Bruijn graph (DBG) in genome assembly^41–43^ prompted us to investigate its potential along this route. Clearly, a DNA strand can be assembled only if the strand path is intact in DBG. We run simulations to estimate the probability of a strand path maintaining its integrity with variant strand copies containing various rates and types of errors. Here, this probability was regarded as the theoretical maximal strand reconstruction rate (***S***_***m***_) of DBG-based strand reconstruction. Strand copies in a range of 3 to 25 were considered, and the detailed simulation results are provided in **Data S1**. As shown by the representative results obtained with five and ten strand copies (**Fig. S1C)**, the ***S***_***m***_ values, as expected, are inversely related to the error rates but are similar regardless of the error types at the same error rates. Importantly, with merely ten sequence copies, the sequence path in DBG shows high robustness even with a high strand error rate of 5%, proving the feasibility of DBG-based strand reconstruction. As illustrated in **Fig. 1B**, we then proposed the DBGPS algorithm for error-free reconstruction of strands, which comprises two stages as follows:

**Fig. 1.**
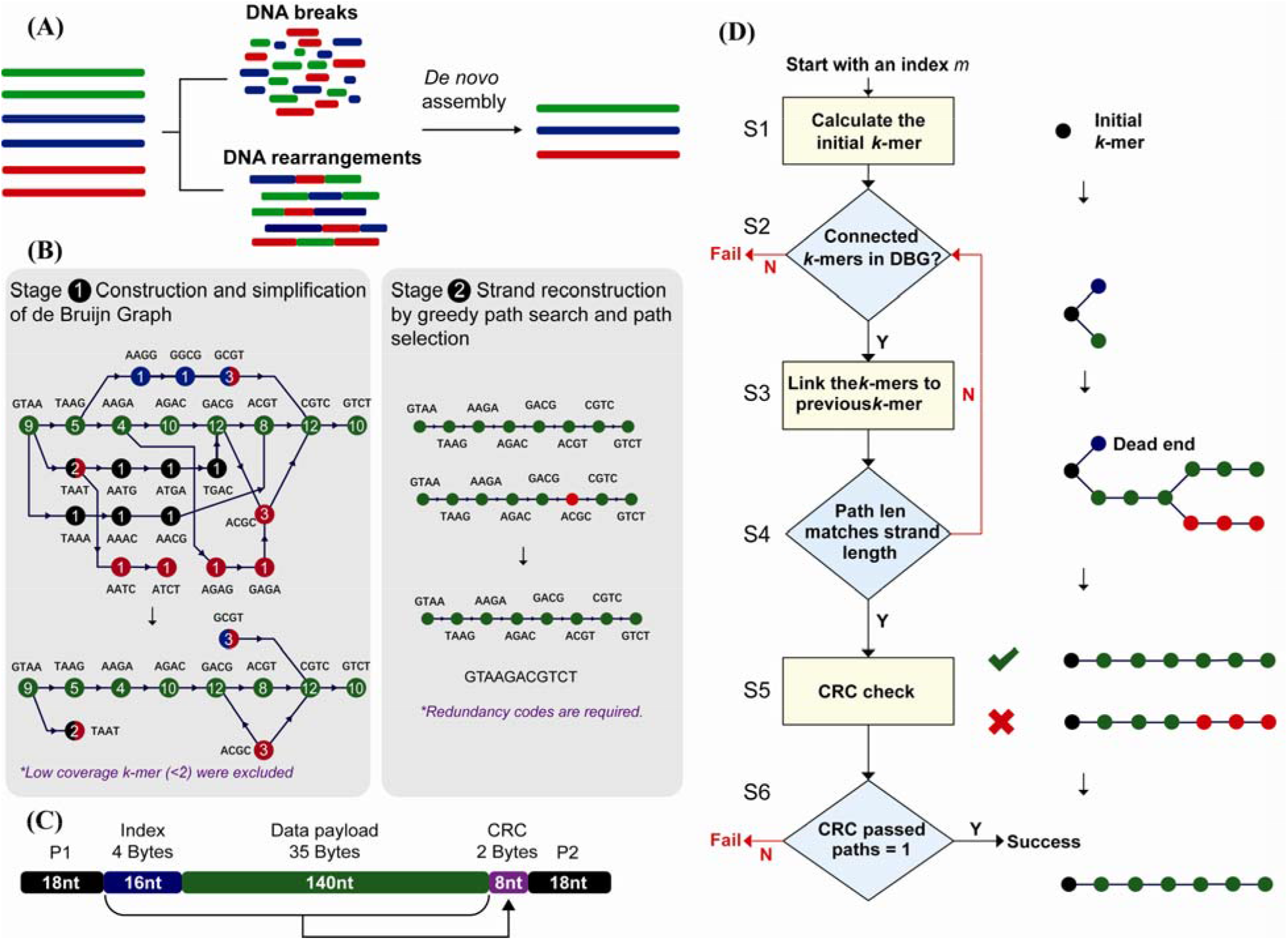
*De novo* assembly-based strand reconstruction for DNA data storage. (A) The issues of DNA breaks and rearrangements in DNA data storage and the proposed *de novo* assembly-based strategy for dealing with them. (B) The two-stage *de novo* assembly process of the proposed de Bruijn graph-based greedy path search algorithm (DBGPS). The representative de Bruijn graph in stage 1 was constructed from the nice error-rich sequence copies shown in **Fig. S1A** with a *k*-mer size of four. The circles stand for the *k*-mer nodes. The numbers inside the circles are the occurrences, *i*.*e*., coverages, of corresponding *k*-mers. The correct sequence is represented by the path of green nodes. (C) The designed strand structure for DBGPS algorithm. (D) The workflow of the greedy path search and path selection process.

#### Stage 1. Construction and simplification of DBG

The construction of DBG here refers to *k*-mer counting^49,50^. The connections between the *k*-mers do not need to be constructed. The *k*-mer counting data in a hash table is well-suited for the greedy path search step in *Stage 2*. Massive errors could cause enormous noise *k*-mers in DBG. The exclusion of noise *k*-mers is crucial for reducing the computing complexity in *Stage 2*. The occurrences of specific errors are low-probability events individually. Therefore, the occurrences, *i*.*e*., coverages, of noise *k-*mers are generally lower than those of the correct ones. As shown by Monte Carlo simulations, coverage is a suitable indicator for the exclusion of noise *k*-mers (**Fig. S2**). The *k*-mer size is an important parameter for strand reconstruction. With a specific *k*-mer size, there is a theoretical upper limit to the volume of data that can be decoded. For example, only a tiny amount of data can be decoded with a *k*-mer size of three. In DBG theory, for each *k*-mer, the front *k*-1 bases were used for positioning, and the terminal base was used for path extension, *i*.*e*., encoding of fresh data. Due to the greedy path search step in *Stage 2*, each front *k*-1 base combination should be used no more than once to avoid path loops and forks. Based on this principle, the decoding capacity was estimated to be around 1 TB (Tera Bytes, 10^12^ Bytes) with a *k*-mer size of 27. The detailed choices of *k*-mer sizes with data volumes ranging from 1 KB (Kilo Bytes, 10^3^ Bytes) to 1 EB (Exa Bytes, 10^18^ Bytes) are listed in **Table S1**. More details of the estimation are provided in the **Supplementary Materials**.

#### Stage 2. Strand reconstruction by greedy path search and path selection

Simply, the strand reconstruction process is achieved by greedy path searching for strand candidates and path selection by the embedded EC codes for the correct strand. To facilitate this process, we designed a strand structure as shown in **Fig. 1C**, which contains 16 nt index, 140 nt data payload, and 8 nt Cyclic Redundancy Check (CRC) code, flanked by landing sites for sequencing primers. As illustrated in **Fig. 1D**, six key steps are required to reconstruct the strand sequence of a specific index *m*: ***S1***. Encode the index *m* into a DNA string and calculate the initial *k*-mer. This initial *k*-mer is also the first terminal *k*-mer. ***S2***. Judgment step. Check if there are connected *k*-mers for each terminal *k*-mer. The terminal *k*-mers without connected *k*-mers are marked as a dead-ends. Strand reconstruction fails if all the terminal *k*-mers are dead-ends. ***S3***. Connect all connected *k*-mers to the corresponding terminal *k*-mers. ***S4***. Step of judgment. Check if the path length matches the strand length. If not, go to *S2*. If so, go to *S5*. ***S5***. Perform parity check for each path candidate using the embedded CRC codes. ***S6***. Step of judgment. Only if exactly one path passes the CRC check, that path is selected as the correct path, *i*.*e*., the strand sequence of index *m*. Strand reconstruction fails if multiple or no paths pass the CRC check. DBGPS will continue to assemble the next strand of index (*m*+1) until all possible indexes are processed.

### The error handling capability and large data scale performance of DBGPS

As shown in **Fig. 2A-E**, we ran simulations to estimate the error handling capability of DBGPS in comparison with the multiple-alignment algorithm (MA) using twenty strand copies. Remarkably, as shown in **Fig. 2AB**, DBGPS shows outstanding performance in handling DNA breaks and rearrangements. In contrast, MA quickly loses its decoding capability with the introduction of a small rate of DNA breaks and rearrangements. DBGPS also shows a clear advantage in handling indels, especially when the error rate is high (**Fig. 2C**). In the case of substitutions (**Fig. 2D**), DBGPS shows a decrease in strand decoding rates when high rates of substitutions are introduced. It has been reported that even with the low-quality synthesis method of light-directed synthesis, the overall error rate was estimated to be around 6%^51^. Thus, the slight deficiency of DBGPS in the handling of high rates of substitutions won’t hamper its practical application in DNA data storage. In practice, the errors are mixtures of different types of errors. We then ran simulations with mixed errors, including substitutions, indels, DNA breaks, and rearrangements in a ratio of 1:1:2:1:1 respectively. As expected, compared with MA, DBGPS shows substantial advantages in the handling of mixed errors as shown in **Fig. 2E**. Next, to investigate how the strand copy number affects the performance of DBGPS and MA, we ran simulations with various strand copies with a fixed error rate of 3% (1.5% substitutions, 0.75% insertions, and 0.75% deletions). As shown in **Fig. 2F**, DBGPS shows a higher *S*_*r*_ value than MA in general, except with extremely low copy numbers under six. The strand reconstruction rates (*S*_*r*_) of DBGPS and MA both increase significantly by introducing more strand copies. With more strand copies introduced, the *S*_*r*_ value of DBGPS increases more quickly and surpasses that of MA at the point of seven.

**Fig. 2.**
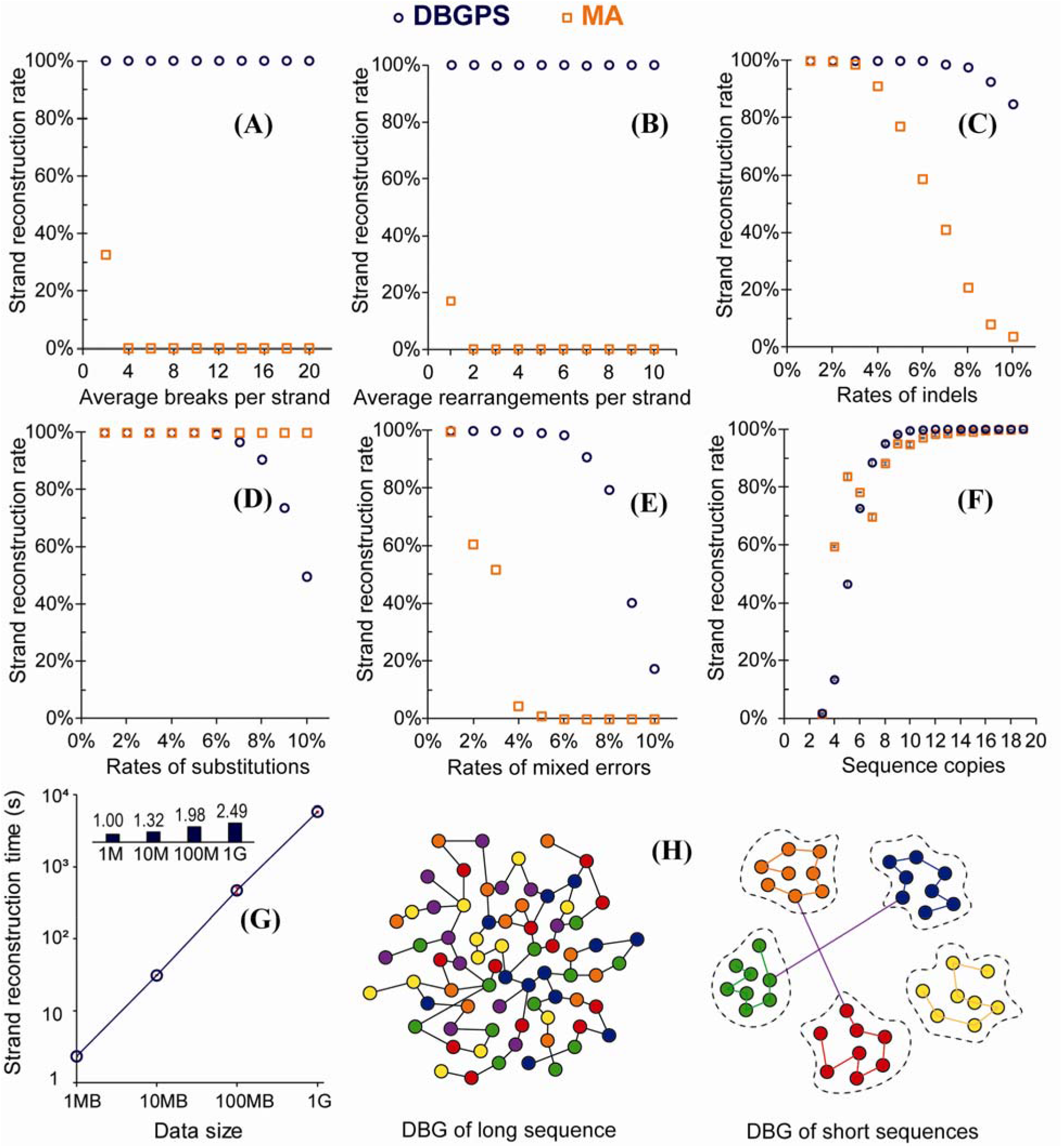
Error-handling capabilities of DBGPS in comparison with MA and large-scale simulations. With twenty sequence copies, the simulated performance of DBGPS and MA in handling various rates of (A) DNA breaks, (B) DNA rearrangements, (C) indels, (D) substitutions, and (E) mixed errors. The mixed errors comprise DNA breaks, DNA rearrangements, substitutions, insertions, and deletions in a ratio of 1:1:2:1:1. (F). Strand reconstruction rates with various strand copies containing 3% error mixtures of substitutions (1.5%), insertions (0.75%), and deletions (0.75%). (G). Strand reconstruction time by DBGPS with data scales ranging from 1 MB to 1 GB. The small bar chart at the top shows the fold changes in reconstruction time per strand compared to the 1 MB scale. (H). Illustration of the differences between the DBG constructed with massive short DNA sequences and that constructed with long DNA sequence(s).

To evaluate the performance of DBGPS with large data volumes, a series of simulations were performed with data sizes ranging from 1 MB to 1 GB. For each data size, three independent simulations were performed using random seeds of 1, 2 and 3 for the generation of the DNA droplets/strands. Preliminary simulations at the GB level showed a significant increase in reconstruction time per strand caused by entanglements of strand paths in DBG, *i*.*e*., repeated presentation of *k*-mers in different strands. To solve this problem, a strand filtering process, as illustrated in **Fig. S3**, was designed and applied to filter out the entangled strands. The DNA strand sequences after filtering were utilized for the generation of error-rich copies. Error-rich strand sequences were simulated with a copy number of 25 and an error rate of 3% (1.5% substitutions, 0.75% insertions, 0.75% deletions). *K*-mer counting has previously been demonstrated to be a linear problem^49,50,52,53^, and this finding was confirmed in this study (**Fig. S4**). For strand reconstruction by DBGPS, the reconstruction time per strand was only slightly increased by scaling up the data size from 1 MB to 1 GB (**Fig. 2G**). A time complexity of *O*(*nlogn*), where *n* stands for data size, was revealed by a fitting experiment (**Fig. S5**). Importantly, no significant change in decoding accuracy was observed with the data volumes ranging from 1 MB to 1 GB (**Fig. S6**). DBG-based sequence reconstruction has been widely studied with the genome assembly problem, revealing the challenge of accurate sequence reconstruction by DBG^41,42,54^. It should be clarified that the sequence reconstruction problem in DNA data storage is very different from that of genome assembly. Rather than long sequences, the DNA sequences in DNA data storage are massive short fragments with a fixed length of 100-300 bp. As illustrated in **Fig. 2H**, the DBG constructed with short fragments shows significant differences from the DBG constructed with long sequence. The nodes in DBG of long sequences are tightly connected. In contrast, the nodes in the DBG of short DNA sequences are spontaneously separated by *k*-mer editing distance. This was proved by analysis of the cross-linked strands at data volumes ranging from 1 MB to 1 GB, as shown in **Fig. S6**. Importantly, these cross-linked strands can be eliminated by the filtering process illustrated in **Fig. S3**, enabling the high efficiency of DBGPS with large datasets. Furthermore, the short fragments in DNA data storage are indexed and are embedded with EC codes, guaranteeing accurate assembly of the original information.

### Experimental verification of the robustness of DBGPS

A 6.8 MB zipped file (6,818,623 bytes, **Data S2**) of ten pictures of Dunhuang murals was encoded into 210,000 DNA strands (**Data S3**) with the structure shown in **Fig. 1C**. The strand redundancy is 7.8%, which supports reliable data recovery with more than 95% of strand sequences. As shown in **Fig. 3A**, the designed oligos were synthesized by Twist Bioscience and the obtained oligos were dissolved in ddH_2_O to generate a “master pool”. Three harsh experiments were performed with this “master pool” to obtain low-quality samples with various rates and types of errors. The low-quality samples obtained were then sequenced on the Illumina sequencing platform. The raw reads generated were directly handled by DBGPS for strand reconstruction followed by decoding using the outer fountain codes.

**Fig. 3.**
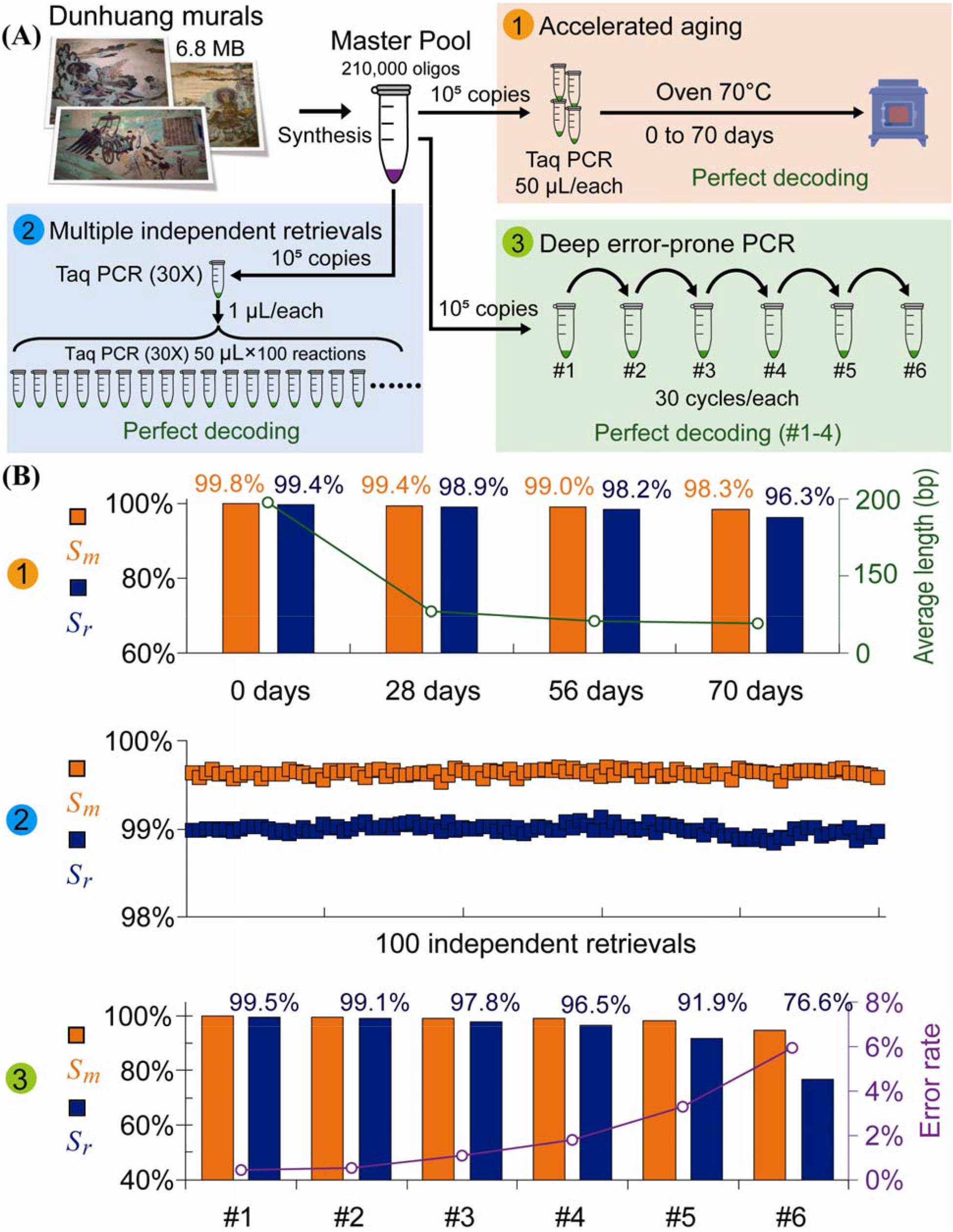
Experimental verification of the robustness of DBGPS. (A) Illustration of the three harsh experiments (1, 2, 3) performed to verify the robustness of DBGPS. A 6.8 MB zipped file of Dunhuang mural pictures was recorded by oligo synthesis, generating an ssDNA “Master Pool” with 210,000 different oligo sequences. *Experiment 1*. Accelerated aging to verify the robustness with DNA degradations (breaks). *Experiment 2*. Multiple data retrievals with intended unspecific amplifications to verify the robustness with strand rearrangements. *Experiment 3*. Deep error-prone PCR to introduce errors. (B) Data retrieval details of the three experiments. All the data retrievals of the three experiments are perfect except ePCR#5 and ePCR#6. The strand reconstruction details by CL-MA are provided in **Table S2**.

#### Robust data retrieval with accelerated aging samples

*In silico* simulation has shown that DBGPS can well handle high rates of DNA breaks. Accelerated aging experiments were performed to further confirm its practical performance in tolerating DNA degradation. The purified PCR products in elution buffer were incubated at 70 °C for a prolonged period of 0, 28, 56, and 70 days. As shown by the Agilent 2100 Bioanalyzer analysis, the integrity of the DNA strands (200bp) has been severely damaged after incubation at 70 °C for merely 28 days (**Fig. S7** and **Fig. S8**). Such a degree of degradation made it impossible to decode the original information using CL-MA (**Table S2**). By contrast, DBGPS achieves high *S*_*r*_ values from all accelerated aging samples, ensuring accurate data retrievals (**Fig. 3B** and **Table S3**). Notably, a high *S*_*r*_ value of 96.3% was even obtained with the sample that was treated at 70 °C for 70 days. Data retrievals from accelerated aged DNAs have been reported^21,35,55^. In these studies, the authors tested the effects of different methods for the preservation of DNA molecules, among which embedment in silicon beads was shown to be the best, allowing recovery of information after treatment at 70 °C for 1 week. According to the authors, this was thermally equivalent to storing information on DNA at 9.4°C for 2,000 years^21^. Based on this estimation, the DBGPS algorithm can retrieve data accurately from DNA solutions preserved at 9.4 °C without particular protection for 20,000 years.

#### Multiple independent data retrievals

Copying and retrieval of data stored in DNAs require PCR-based amplification. As a typical biological process, unspecific amplification occurs occasionally, leading to DNA rearrangements. To demonstrate the effectiveness of DBGPS in the handling of DNA rearrangements, 100 independent data retrievals with PCR-amplified products were performed (**Fig. 3A**), in which unspecific amplifications were intendedly introduced. The CL-MA based decoding shows low *S*_*r*_ values of less than 12% with three representative samples (**Table S2**). In contrast, high *S*_*r*_ values ranging from 98.8% to 99.1% were obtained with DBGPS, ensuring perfect data recovery in all retrievals (**Fig. 3B, Data S4**).

#### Data retrieval with deep, error-prone PCR products

A series of six error-prone PCR (ePCR) amplifications were performed to introduce large numbers of base errors at various rates. All ePCR reactions were performed for 30 thermal cycles. Regardless of rough amplification conditions expected to introduce a large number of errors, strikingly high *S*_*r*_ values in a range of 96.6% to 99.5% were obtained for ePCR#1-4, ensuring accurate data recovery (**Table S4**). Decrease in *S*_*r*_ was observed as expected with increased cycles of ePCR. More PCR errors were expected to increase branch paths, raising the necessary cutoff for noise *k*-mer elimination. Consequently, more correct *k*-mers were consequently discarded by mistake, resulting in lower *S*_*r*_ values. Although the data retrievals with ePCR#5 and ePCR#6 failed, the total 120 cycles of amplification performed in ePCR#1-4 can already guarantee sufficient reliable data copies.

## Discussion

DNA data storage technology maintains the order of information by defined sequences of nucleic acid chains. Different from the traditional plane medium, which requires a surface to maintain the information order, the nucleic acid chains can be detached from the writing surface and distributed in a three-dimensional space without affecting the data integrity. This gives DNA data storage a huge increase in data density. However, the chain issues of breakages and incorrect linkages, *i*.*e*., DNA breaks and rearrangements, need to be handled. As illustrated in **Fig. S9**, in a radical view, substitutions and indels can also be regarded as special cases of “DNA rearrangements”, implying the importance of handling these fundamental chain issues. However, these issues are difficult to tackle using traditional EC codes. In this study, we developed DBGPS, a *de novo* assembly-based strand reconstruction algorithm based on DBG theory, which can take advantage of the unique “multicopy” feature for the error-free reconstruction of strand sequences. We verified the effectiveness of DBGPS in the handling of DNA breaks and rearrangements, as well as substitutions and indels, by both dry and wet experiments. With this algorithm, we were able to accurately and rapidly reconstruct DNA strands from error-rich strand copies that were mixed together without a clustering step.

The most important contribution of this work is the handling of data reliability issues caused by DNA degradation. The importance of handling DNA degradation in DNA data storage has been well discussed in a recent review^36^. Different from the previous studies, which focused on protecting the DNA molecules from corruption^21,35,36,55^, here we focused on the implementation of a computational process to reconstruct the strand sequences even if the strands are broken into small fragments of dozens of bases. As proved by the accelerated aging experiment, we can well handle the data robustness issue caused by DNA degradation with DBGPS. It is worth noting that the encoded information from a DNA solution that has been incubated at 70 °C for 70 days without any particular protection can be accurately recovered with our algorithm. The degree of damage to DNA under this heating condition was estimated to be equivalent to that under 9.4 °C for more than 20,000 years^21^. This period is already longer than the oldest written record of human civilization, the Cuneiform, which dates back about 5,500 years ago^56^. Such unprecedented robustness highlights the importance of our method and suggests the great potential of our DBGPS algorithm in DNA data storage. DBGPS is presumably compatible with a variety of DNA preservation methods, *e*.*g*., silicon beads^21^, nanoparticles^55^ and alkaline salts^35^, to further enhance data stability. Altogether, it is reasonable to believe that DNA data storage technology might enable us to preserve human culture for a long period of time, even after the unavoidable destruction of mankind in the distant future. Another significant contribution of this study is the efficient handling of unspecific amplification, which improves the robustness of DNA data storage significantly. Retrieval of DNA data requires PCR-based amplification of strands. As a typical biological process, PCR is stochastic in nature. This stochastic feature frequently leads to unspecific amplification, threatening the robustness of DNA data storage. With DBGPS, we achieved robust data retrieval even if unspecific amplification occurs, as proved by the 100 independent retrievals with intentionally introduced unspecific amplification.

Based on traditional information theory, redundancy data in the form of “repetition” is unfavorable since the production of additional data copies on traditional media is cost-ineffective and time-consuming. By contrast, data repetition in the form of DNA strand copies can be obtained inexpensively and speedily. Thus, the authors believe that the traditional definition of “coding efficiency” needs to be adapted to the DNA data storage channel. With current technologies, producing a single copy of DNA data is not even doable. It has been reported that serial dilutions to reduce the copy numbers of information-bearing DNA strands till the average copy number is below ten result in massive strand dropouts, making reliable data retrieval impractical^46^. Even if a single copy of DNA data is obtainable, it may be damaged due to DNA degradation under certain conditions or during long-term storage. Taken together, the design for DNA storage should be based on reality and necessity, allowing for the presence of more than one copy of DNA strands and not too many copies to maintain high information density. In other words, “multicopy” is crucial for the robustness of DNA data storage and should ideally be maintained in a reasonable range. Data robustness, physical density, and economy together determine the technical advancements of a storage medium. By taking advantage of the “multicopy” feature, this work significantly improved the robustness of DNA data storage while maintaining high physical and logical density. The latter is an important indicator highly related to writing costs. As detailed in **Table 1**, we achieved a physical density of 295 PB/g and a logical density of 1.30 bits/cycle at a data scale of 6.8 MB. Recently, Anavy *et al*. have reported a logical density of 1.53 bits/cycle using composite DNA letters at a scale of 6.4 MB. However, a low physical density of 5.9 PB/g is reported due to the high coverage required by composite DNA letters^57^. Compared to the study of Anavy *et al*., we achieve much higher physical density and data robustness at the cost of a slight decrease in logical density. It’s worth mentioning that we obtained a strand dropout rate of 1.79% at a scale of 6.8 MB, which is significantly lower than the reported dropout rate of 3.60% at a scale of 2.14 MB by Erlich *et al*., revealing the technical advances of DBGPS.

**Table 1.** Key achievements of this work in comparison with prior DNA storage studies. Note: primers were considered in the calculation of logical density (bits per synthesis cycle). The long-term stability was estimated based on the study by Grass *et al*.^21^.

|  | **Church**<br>***et al.*<sup>3</sup>** | **Goldman**<br>***et al.*<sup>24</sup>** | **Grass**<br>***et al.*<sup>21</sup>** | **Erlich**<br>***et al.*<sup>22</sup>** | **Organick**<br>***et al.*<sup>23</sup>** | **Leon**<br>***et al.*<sup>57</sup>** | **Antkowi**<br>**aket *al.*<sup>26</sup>** | **This work** |
| --- | --- | --- | --- | --- | --- | --- | --- | --- |
| **Data size**<br>**(MB)** | 0.53 | 0.74 | 0.08 | 2.14 | 200.2 | 6.42 | 0.1 | **6.8** |
| **Total oligos** | 54,898 | 153,335 | 4,991 | 72,000 | 13,400,000 | 217,000 | 16,383 | **210,000** |
| **Logical**<br>**density**<br>**(Bits/cycle)** | 0.6 | 0.19 | 0.86 | 1.19 | 0.81 | 1.52 | 0.94 | **1.30** |
| **Physical**<br>**density**<br>**(PB/g)** | NA | NA | NA | 215 | NA | 5.9 | NA | **295** |
| **Long-term**<br>**stability**<br>**(9.4°C)** | - | - | ~2,000<br>years | - | - | - | - | **~20,000**<br>**years** |

Although the mechanism of low coverage *k*-mer exclusion works well with most of the data retrievals in this study, we observed a significant decrease in strand reconstruction rate when more and more errors are introduced (**Fig. 3B, Table S4**). Future incorporation of redundancy codes that can help to identify and exclude the noise *k*-mers is expected to further improve the efficiency of DBGPS, which is particularly important in the extreme case of high error rates^26,51^. The recent study by Antkowiak *et al*. revealed the potential of low-quality synthesis methods, *e*.*g*. photolithographic and electrochemical synthesis, in reducing the writing cost of DNA data storage. However, due to the high synthesis error rate, a low strand recovery rate of 83% was reported using the CL-MA method^26^. Interestingly, DBGPS combined with a simple mechanism of inserting one error-checking base every few bases (block checking codes)^58^ could be a potential solution to this problem. With block lengths ranging from 3 to 10, simulation tests of this strategy on the sequencing data from the study of Antkowiak *et al*.^26^ revealed high strand reconstruction rates ranging from 98.3% to 99.6% (**Table S6**), suggesting a potential strategy in low-cost, inaccurate DNA synthesis technologies for data storage.

## Acknowledgments

This work was supported by National Natural Science Foundation of China under the grants 21621004 to YJY, Seed Foundation of Tianjin University to LFS, Natural Science Foundation of Shandong Province under the grants ZR2017LC006 to FG. The authors would like to thank Dr. Yan Zhang for his great efforts and constructive comments which have helped to improve this manuscript significantly. The authors also would like to thank Dr. Feng Gao, Dr. Hao Qi, Dr. Weigang Chen, Dr. Yu Lin, and Dr. Sheng Ye for their insightful discussions.

## Author contributions

LFS and YJY conceived the study. LFS proposed and designed the algorithm, participated in the experiment design, and data analysis. FG performed the performance testing and participated in the algorithm and data analysis. ZYG performed all the experiments. CYG and RX participated in the implementation of DBGPS-C. JJT and XC participated in the decoding complexity analysis. LBZ, MZH and JYX helped with the experiments. BZL and YJY edited the manuscript and supervised the whole work.

## Competing interests

A patent covering the encoding structure and the decoding process has been filed (CN 110190858 A).

## Data and materials availability

All the original algorithms proposed in this study were implemented with Python and the source codes are available on GitHub at URL: https://github.com/Scilence2022/DBGPS_Python. A compiled C implementation of DBGPS is available at https://github.com/Scilence2022/DBGPS-C. The source codes of the C version can be obtained for academic usage upon request to the authors. All relevant data can be retrieved at: https://figshare.com/projects/DBGPS/123871.

## Notes

https://switch-codes.coding.net/public/switch-codes/DNA-De-Bruijn-Decoding/git/files

### Competing Interest Statement

Li-Fu Song, Ying-Jin Yuan and Feng Geng have filed a patent covering the encoding structure and the de Bruijn graph based decoding process (CN 110190858 A).

### Summary of Updates

Data scale: 0.4 MB -> 6.8 MB; C verision of DBGPS with multi-threats k-mer counter;

## References

1. van der Valk, T. et al. Million-year-old DNA sheds light on the genomic history of mammoths. Nature 591, 265–269; 10.1038/s41586-021-03224-9 (2021).

2. Zhirnov, V., Zadegan, R. M., Sandhu, G. S., Church, G. M. & Hughes, W. L. Nucleic acid memory. Nature materials 15, 366–370; 10.1038/nmat4594 (2016).

3. Church, G. M., Gao, Y. & Kosuri, S. Next-generation digital information storage in DNA. Science (New York, N.Y.) 337, 1628; 10.1126/science.1226355 (2012).

4. Ceze, L., Nivala, J. & Strauss, K. Molecular digital data storage using DNA. Nature reviews. Genetics 20, 456–466; 10.1038/s41576-019-0125-3 (2019).

5. Ping, Z. et al. Carbon-based archiving: current progress and future prospects of DNA-based data storage. GigaScience 8; 10.1093/gigascience/giz075 (2019).

6. Chen, W. et al. An artificial chromosome for data storage. National Science Review; 10.1093/nsr/nwab028 (2021).

7. Tabatabaei, S. K. et al. DNA punch cards for storing data on native DNA sequences via enzymatic nicking. Nature communications 11, 1742; 10.1038/s41467-020-15588-z (2020).

8. Koch, J. et al. A DNA-of-things storage architecture to create materials with embedded memory. Nature biotechnology 38, 39–43; 10.1038/s41587-019-0356-z (2020).

9. Lu, X. & Ellis, T. Self-replicating digital data storage with synthetic chromosomes. National Science Review 8, 1763; 10.1093/nsr/nwab086 (2021).

10. Meiser, L. C. et al. Reading and writing digital data in DNA. Nature protocols 15, 86–101; 10.1038/s41596-019-0244-5 (2020).

11. Song, L., Deng, Z., Gong, Z., Li, L. & Li, B. Large-Scale de novo Oligonucleotide Synthesis for Whole-Genome Synthesis and Data Storage: Challenges and Opportunities. Frontiers in bioengineering and biotechnology 9, 689797; 10.3389/fbioe.2021.689797 (2021).

12. Chandak, S. et al. Overcoming High Nanopore Basecaller Error Rates for DNA Storage via Basecaller-Decoder Integration and Convolutional Codes. In ICASSP 2020 - 2020 IEEE International Conference on Acoustics, Speech and Signal Processing (ICASSP) (IEEE Monday, May 4, 2020 - Friday, May 8, 2020), pp. 8822–8826.

13. Press, W. H., Hawkins, J. A., Jones, S. K., Schaub, J. M. & Finkelstein, I. J. HEDGES error-correcting code for DNA storage corrects indels and allows sequence constraints. Proceedings of the National Academy of Sciences of the United States of America 117, 18489–18496; 10.1073/pnas.2004821117 (2020).

14. Dong, Y., Sun, F., Ping, Z., Ouyang, Q. & Qian, L. DNA storage: research landscape and future prospects. National Science Review; 10.1093/nsr/nwaa007 (2020).

15. Xu, C., Zhao, C., Ma, B. & Liu, H. Uncertainties in synthetic DNA-based data storage. Nucleic acids research 49, 5451–5469; 10.1093/NAR/GKAB230 (2021).

16. Lee, H. et al. Photon-directed multiplexed enzymatic DNA synthesis for molecular digital data storage. Nature communications 11, 5246; 10.1038/s41467-020-18681-5 (2020).

17. Banal, J. L. et al. Random access DNA memory using Boolean search in an archival file storage system. Nature materials 20, 1272–1280; 10.1038/s41563-021-01021-3 (2021).

18. Lin, K. N., Volkel, K., Tuck, J. M. & Keung, A. J. Dynamic and scalable DNA-based information storage. Nature communications 11, 2981; 10.1038/s41467-020-16797-2 (2020).

19. Lee, H. H., Kalhor, R., Goela, N., Bolot, J. & Church, G. M. Terminator-free template-independent enzymatic DNA synthesis for digital information storage. Nature communications 10, 2383; 10.1038/s41467-019-10258-1 (2019).

20. Bancroft, C., Bowler, T., Bloom, B. & Clelland, C. T. Long-term storage of information in DNA. Science (New York, N.Y.) 293, 1763–1765; 10.1126/science.293.5536.1763c (2001).

21. Grass, R. N., Heckel, R., Puddu, M., Paunescu, D. & Stark, W. J. Robust chemical preservation of digital information on DNA in silica with error-correcting codes. Angewandte Chemie (International ed. in English) 54, 2552–2555; 10.1002/anie.201411378 (2015).

22. Erlich, Y. & Zielinski, D. DNA Fountain enables a robust and efficient storage architecture. Science (New York, N.Y.) 355, 950–954; 10.1126/science.aaj2038 (2017).

23. Organick, L. et al. Random access in large-scale DNA data storage. Nature biotechnology 36, 242–248; 10.1038/nbt.4079 (2018).

24. Goldman, N. et al. Towards practical, high-capacity, low-maintenance information storage in synthesized DNA. Nature 494, 77–80; 10.1038/nature11875 (2013).

25. Gao, Y., Chen, X., Qiao, H., Ke, Y. & Qi, H. Low-Bias Manipulation of DNA Oligo Pool for Robust Data Storage. ACS synthetic biology 9, 3344–3352; 10.1021/acssynbio.0c00419 (2020).

26. Antkowiak, P. L. et al. Low cost DNA data storage using photolithographic synthesis and advanced information reconstruction and error correction. Nature communications 11, 5345; 10.1038/s41467-020-19148-3 (2020).

27. Cyrus Rashtchian et al. Clustering Billions of Reads for DNA Data Storage 30 (2017).

28. Levenshtein, V. I. Efficient reconstruction of sequences. IEEE Trans. Inform. Theory 47, 2–22; 10.1109/18.904499 (2001).

29. Bhardwaj, V., Pevzner, P. A., Rashtchian, C. & Safonova, Y. Trace Reconstruction Problems in Computational Biology. IEEE Trans. Inform. Theory 67, 3295–3314; 10.1109/tit.2020.3030569 (2021).

30. Sabary, O., Yucovich, A., Shapira, G. & Yaakobi, E. Reconstruction Algorithms for DNA-Storage Systems (2020).

31. Shomorony, I. & Heckel, R. DNA-Based Storage: Models and Fundamental Limits. IEEE Trans. Inform. Theory 67, 3675–3689; 10.1109/TIT.2021.3058966 (2021).

32. Cheraghchi, M., Gabrys, R., Milenkovic, O. & Ribeiro, J. Coded Trace Reconstruction. IEEE Trans. Inform. Theory 66, 6084–6103; 10.1109/TIT.2020.2996377 (2020).

33. Kiah, H. M., Puleo, G. J. & Milenkovic, O. Codes for DNA Sequence Profiles. IEEE Trans. Inform. Theory 62, 3125–3146; 10.1109/TIT.2016.2555321 (2016).

34. Batu, T., Kannan, S., Khanna, S. & Mcgregor, A. Reconstructing Strings from Random Traces. Proceedings of the Annual ACM-SIAM Symposium on Discrete Algorithms 15; 10.1145/982792.982929 (2004).

35. Kohll, A. X. et al. Stabilizing synthetic DNA for long-term data storage with earth alkaline salts. Chemical communications (Cambridge, England) 56, 3613–3616; 10.1039/d0cc00222d (2020).

36. Matange, K., Tuck, J. M. & Keung, A. J. DNA stability: a central design consideration for DNA data storage systems. Nature communications 12, 1358; 10.1038/S41467-021-21587-5 (2021).

37. Zorita, E., Cuscó, P. & Filion, G. J. Starcode: sequence clustering based on all-pairs search. Bioinformatics (Oxford, England) 31, 1913–1919; 10.1093/bioinformatics/btv053 (2015).

38. Edgar, R. C. MUSCLE: multiple sequence alignment with high accuracy and high throughput. Nucleic acids research 32, 1792–1797; 10.1093/nar/gkh340 (2004).

39. Katoh, K., Misawa, K., Kuma, K.-i. & Miyata, T. MAFFT: a novel method for rapid multiple sequence alignment based on fast Fourier transform. Nucleic acids research 30, 3059–3066; 10.1093/nar/gkf436 (2002).

40. Raphael, B., Zhi, D., Tang, H. & Pevzner, P. A novel method for multiple alignment of sequences with repeated and shuffled elements. Genome research 14, 2336–2346; 10.1101/gr.2657504 (2004).

41. Zerbino, D. R. & Birney, E. Velvet: algorithms for de novo short read assembly using de Bruijn graphs. Genome research 18, 821–829; 10.1101/gr.074492.107 (2008).

42. Pevzner, P. A., Tang, H. & Waterman, M. S. An Eulerian path approach to DNA fragment assembly. Proceedings of the National Academy of Sciences of the United States of America 98, 9748–9753; 10.1073/pnas.171285098 (2001).

43. N. G. de Bruijn. A combinatorial problem. Indagationes Mathematicae 49, 758–764 (1946).

44. Bankevich, A., Bzikadze, A. V., Kolmogorov, M., Antipov, D. & Pevzner, P. A. Multiplex de Bruijn graphs enable genome assembly from long, high-fidelity reads. Nature biotechnology; 10.1038/s41587-022-01220-6 (2022).

45. Organick, L. et al. Experimental Assessment of PCR Specificity and Copy Number for Reliable Data Retrieval in DNA Storage (2019).

46. Chen, Y.-J. et al. Quantifying molecular bias in DNA data storage. Nature communications 11, 3264; 10.1038/s41467-020-16958-3 (2020).

47. Heckel, R., Mikutis, G. & Grass, R. N. A Characterization of the DNA Data Storage Channel. Scientific reports 9, 9663; 10.1038/s41598-019-45832-6 (2019).

48. Hao, M. et al. A mixed culture of bacterial cells enables an economic DNA storage on a large scale. Communications biology 3, 416; 10.1038/s42003-020-01141-7 (2020).

49. Pandey, P., Bender, M. A., Johnson, R., Patro, R. & Berger, B. Squeakr: an exact and approximate k-mer counting system. Bioinformatics (Oxford, England) 34, 568–575; 10.1093/bioinformatics/btx636 (2018).

50. Marçais, G. & Kingsford, C. A fast, lock-free approach for efficient parallel counting of occurrences of k-mers. Bioinformatics (Oxford, England) 27, 764–770; 10.1093/bioinformatics/btr011 (2011).

51. Lietard, J. et al. Chemical and photochemical error rates in light-directed synthesis of complex DNA libraries. Nucleic acids research 49, 6687–6701; 10.1093/nar/gkab505 (2021).

52. Melsted, P. & Pritchard, J. K. Efficient counting of k-mers in DNA sequences using a bloom filter. BMC bioinformatics 12, 333; 10.1186/1471-2105-12-333 (2011).

53. Heng Li. Fast and simple k-mer counters. Available at https://github.com/lh3/kmer-cnt.

54. Compeau, P. E. C., Pevzner, P. A. & Tesler, G. How to apply de Bruijn graphs to genome assembly. Nature biotechnology 29, 987–991; 10.1038/nbt.2023 (2011).

55. Chen, W. D. et al. Combining Data Longevity with High Storage Capacity— Layer□by□Layer DNA Encapsulated in Magnetic Nanoparticles. Adv. Funct. Mater. 29, 1901672; 10.1002/adfm.201901672 (2019).

56. Walker, C. B. F. Cuneiform (University of California Press; British Museum, Berkeley CA, London, 1987).

57. Anavy, L., Vaknin, I., Atar, O., Amit, R. & Yakhini, Z. Data storage in DNA with fewer synthesis cycles using composite DNA letters. Nature biotechnology; 10.1038/s41587-019-0240-x (2019).

58. Song, L. & Zeng, A.-P. Orthogonal Information Encoding in Living Cells with High Error-Tolerance, Safety, and Fidelity. ACS synthetic biology 7, 866–874; 10.1021/acssynbio.7b00382 (2018).

