## Supplemental info. for "Robust retrieval of data stored in DNA by de Bruijn graph-based *de novo* strand assembly"

**nature**

**communications**

Supplementary Materials for

Robust data storage in DNA by *de novo* assembly-based strand reconstruction

Lifu Song^1,2^, Feng Geng^3^, Ziyi Gong^1,2^, Xin Chen^4^, Jijun Tang^5,7^, Chunye Gong^6^, Libang Zhou^8^, Rui Xia^6^, Mingzhe Han^1,2^, Jingyi Xu^1,2^, Bingzhi Li^1,2^*, Yingjin Yuan^1,2^*

**This file includes:**

Materials and Methods

Figs. S1 to S12

Tables S1 to S6

Captions for Movies S1

Captions for Data S1 to S4

References

**Other Supplementary Materials for this manuscript include the following:**

Movie S1

Data S1 to S4

Materials and Methods

**Library design, preparation of PCR samples, and sequencing**

A zipped file (6.8 M, **Data S1**) of Dunhuang Murals was used as input for library design. A DNA library of 210,000 oligonucleotides with a length of 200 nt was produced by DNA fountain codes (**Data S2**). Each includes 16 nt index, 140 nt data payload, 8 nt CRC codes, and 18 nt ×2 primers: P1 5'-CCTGCAGAGTAGCATGTC-3', P2 5'-CGGATGCATCAGTGTCAG-3'. The oligo pool was synthesized by Twist Bioscience. The synthesized pool was resuspended in ddH_2_O for a final concentration of 34 ng/µL.

All error-prone PCR were performed with Controlled Error-prone PCR Kit, TIANDZ, Beijing China, CAT#:160903-100. Thermo cycle parameters were 94°C for 3 mins, 94°C for 1 min, 45°C for 1 min, 72°C for 30 s, 30 cycles. Six serial error-prone PCR (ePCR) were performed to introduce high rates of errors. The first round ePCR utilized 0.6 µL 10^-1^ diluted samples (~10^5^ copies) of the master pool as templates. The five serial ePCR use 1 µL products from the previous round of ePCR as templates.

For the 100 multiple independent retrievals, we first performed one round of PCR amplification with a total volume of 100 µL using 0.6 µL 10^-1^ diluted master pool solution as templates. Then, 100 independent PCR amplifications were performed, each using 1 µL of the PCR reaction mixture of first round as templates. All the amplifications were performed with Vazyme 2X Rapid Taq Master Mix (CAT#: P222-AA).

For the accelerated aging experiments, ten parallel PCR reactions were performed in a volume of 100 µL with Vazyme 2X Rapid Taq Master Mix (CAT#: P222-AA). For each reaction, 1 µL of diluted master pool solution (~10^5^ molecular copies/µL) was used as templates. Thermo cycle parameters were 95°C for 30 s, 95°C for 15s, 51°C for 15s, 72°C for 6s, for a total of 30 cycles. The obtained PCR products were purified together with the SparkJade DNA purification kit (CAT#:AE0301-B) and eluted with elution buffer. The obtained ~400 µL solution was diluted with 200 µL of elution buffer. The diluted solution was divided into seven 2 mL tubes with screw caps, each holding 50 µL. The tubes were then incubated at 70°C for 28, 56, and 70 days. The tubes were collected after incubation and stored at -20°C until sequencing.

All the PCR products were sequenced by Tianjin Novogene Sequencing Center & Clinical Lab. Sequencing libraries were generated with purified PCR products using the Illumina TruSeq DNA PCR-Free Library Preparation Kit (Illumina, USA) following the manufacturer’s recommendations. A DNase digestion step is applied to break the long strands into small fragments of 200-300 bp if necessary. After digestion, all fragments are collected for further library construction. The library quality was assessed on the Qubit@2.0 Fluorometer (Thermo Scientific) and the Agilent Bioanalyzer 2100 system. All the libraries were sequenced on the Illumina HiSeq platform, and 150 bp paired-end reads were generated.

**DBGPS implementations in Python and C**

A python version of DBGPS was implemented and the source codes are available at https://github.com/Scilence2022/DBGPS_Python. This python implementation used a simple hash dictionary structure for *k*-mer counting. This implementation is not efficient but versatile for testing new ideas with small data volumes. We also implemented a C version of DBGPS which integrates a multi-threads *k*-mer counter from <https://github.com/lh3/kmer-cnt/blob/master/kc-c4.c>. The C version is more than ten times faster than the Python version generally and is much more memory efficient. Several tools were also provided which can be useful for DNA data storage studies. DBGPS-SmKnd is a tool for the calculation of the *K_n_*, *K_d_*, and *S_m_* values of specific sequencing results. DBGPS-ft is a tool for filtering out the entangled strands. We provided the compiled program at <https://github.com/Scilence2022/DBGPS-C>. For academic usage, the source codes can be obtained upon request to the authors.

The default parameters are optimized for decoding the strands encoding the 6.8 MB zipped file in this study. The usage:

```
Usage: DBGPS [options] <input file>
                       [Supporting formats: *fq, *fa, *fq.gz, *fa.gz]
Options:
  -k INT     k-mer size [31]
  -i INT     length of index [16] bp
  -l INT     data encoding length [140] bp
  -t INT     number of threads [3]
  -c INT     minimal k-mer coverage for strand decoding [1]
  -a INT     Initial index [101010102]
  -b INT     End index [101295684]
```

**The DNA Fountain Codes in Python**

The DNA fountain codes used in this study are publicly available at https://github.com/Scilence2022/DBGPS_Python. DNAFountain and DNADroplet objects were implemented to deal with the transcoding between binary and DNA strings. The degree function is re-designed to follow a slightly modified version of robust distribution. In this modification, the probability of degree one droplets is multiplied by ten times. The assignment mechanism of the random degrees and chunk indexes of the Droplets was also modified to support DBGPS based strand reconstruction. In more detail, a random degree table was pre-generated using the modified degree generation function. For the production of DNA Droplets, a specific index is assigned and used for picking a degree number from the pre-generated degree table for each droplet. This index is also used as a seed for the generation of the random data chunk combinations for each droplet. The original implementation of the Glass object is very slow with large files. The decoding function has been overwritten to improve its performance. For generation of the 210,000 DNA strands encoding the 6.8 MB of data, we provided a specialized script “*Produce_6.8M_210K_Droplets.py*”. Python implementation of DBGPS algorithm is provided in the script “*deBruijnGraph.py*”.

Script for encoding of digital data into strand sequences: *encode.py*

```
Usage:
      python encode.py -i input_file -n number_of_droplets -o output.fasta [Options]
Options:
      -h, --help                              Show help information
      -i, --input   <input file>              Input file
      -o, --output  <output file>             Output file
      -r, --redundancy_rate  <number>         Redundancy rate, default 0.05
      -n, --droplet_num  <number>             Number of droplets, default 210,000
      -c, --chunk_size  <size>                Chunk size, default 35 (bytes)
      -s, --seed    <seed>                    Fountain random seed, default 1
      -l, --initial_index  <initial index>    Initial index, default 1
      --index_bytes  <number>                 Length of index codes, default 4 (bytes)
 --ec_bytes  <number>                    Length of ec codes, default 2 (bytes)
```

Script for decoding of original data from strand sequences without primers *decode_DBGPS.py*. The strand sequences need to be constructed by the C version of DBGPS. The usage of this script:

```
Usage:
      python decode_DBGPS.py -i input_file -o outfile [Options]
Options:
      -h, --help                              Show help information
      -i, --input   <input file>              The decoded strands by DBGPS
      -o, --output  <output file>             Output file
      -d, --chunk_size  <size>                Chunk size, default = 35 (bytes)
      -n, --chunk_num  <number>               Chunk number, default = 194,818
      --seed    <seed>                        Fountain random seed, default 1
      --index_bytes  <number>                 Bytes of index codes, default = 4
 --ec_bytes  <number>                    Bytes of ec codes, default = 2
```

Script for decoding of original data from raw sequencing reads: *decode.py.* This script used the Python implementation of DBGPS. It should be noticed that this script is more than ten times slower than the C version of DBGPS in general. The usage of *decode.py*:

```
Usage:
      python decode.py -i input_file -t type_of_seqs -o outfile [Options]
Options:
      -h, --help                              Show help information
      -i, --input   <input file>              Input file
      -t, --file_type   <file type>           Input file type: FastQ, Fasta or Jellyfish dumped k-mers (default)
      -o, --output  <output file>             Output file
      -k, --kmer_size  <number>               k-mer size, default = 21
      -c, --chunk_size  <size>                Chunk size, default = 35 (bytes)
      -n, --chunk_num  <number>               Chunk number, default = 194,818

      -s, --seed    <seed>                    Fountain random seed, default 1
      --cut  <number>                         Cutoff for exclusion of noisy k-mers default=0
      --min_index  <initial index>            Initial index, default = 101010102
      --max_index  <max index>                Max index, default = 101295684
      --index_bytes  <number>                 Length of index codes, default=4 (bytes)
      --ec_bytes  <number>                    Length of ec codes, default = 2 (bytes)
```

**Strand filtering process to avoid entangled strands in DBG**

The filter process is basically a special *k*-mer counting process. As illustrated in **Fig. S3**, when the filter receives a new strand sequence, the highest occurrence/coverage of all the *k*-mers of this sequence is calculated by querying the *k*-mer hash table. Only if the highest coverage is lower than a specific value (a parameter of filtering), the *k*-mers are then inserted into the *k*-mer hash table. Otherwise, this strand is marked as a strand to be dropped out to avoid entanglements of strands. To clarify, the *k*-mer size here should be smaller than the *k*-mer size during decoding by DBGPS. The error rate should be considered with the *k*-mer size setting for strand filtering. If the error rate after noise exclusion is high, a smaller *k*-mer size for strand filtering is preferred.

**Integration of DBGPS with outer erasure codes**

DBGPS is designed as an inner decoding mechanism for the error-free reconstruction of short DNA strands for DNA data storage. It can be easily combined with an outer erasure code, *e.g.* fountain codes or RS codes. It should be noticed that Fountain code is a better choice than RS codes because a strand filtering process can be easily integrated into the encoding stage of fountain codes to filter out the entangled strands that are tricky to handle by DBGPS. For proof of concept, Python implementation of DBGPS using fountain codes as outer codes is available at <https://github.com/Scilence2022/DBGPS_Python>. For large-scale data storage over 1 GB, a strand filtering process as illustrated in **Fig. S3** is highly recommended. A compiled strand filter (DBGPS-ft) is provided at <https://github.com/Scilence2022/DBGPS-C>. As shown in **Fig. S10**, the integrated decoding process is simple. At first, the raw sequencing reads are handled by DBGPS to reconstruct the strand sequences. Then, the reconstructed strands are treated by the outer erasure codes to decode the original data. An end-to-end presentation of the decoding process by DBGPS and outer fountain codes was provided in **Movie S1**. A demo of DBGPS in Ubuntu virtual machine is available at: <https://doi.org/10.6084/m9.figshare.17161742.v3>.

**Choice of *k*-mer size**

In DBG theory, for each *k*-mer, the front *k*-1 bases were used for positioning, and the terminal base was used for path extension, *i.e.* encoding of fresh data. Due to the greedy path search step in *Stage 2*, each front *k*-1 base combination should be used no more than once to avoid path loops/forks. Thus, we estimated the decoding capacity ($D$) of specific size *k*-mers as follows:

$D=4^{k-1}\times2 bits$	(1)

Where $4^{k-1}$ is the number of possible *k*-1 base combinations, 2 bits stands for the encoding capacity of the single base at the 3’-terminal. Based on formula (1), we can obtain the formula for calculation of *k*-mer size with specific data volume *D* in bits:

$k'=\frac{\ln D- \ln2}{\ln4} +1$	(2)

In practice, however, the random errors can also possibly introduce path loops/forks. Thus, the *k*-mer editing distance should be large enough to avoid the formation of forks/loops by the errors. To ensure sufficient *k*-mer editing distance between arbitrary two *k*-mers in DBG, the *k*-mer combination space should be larger enough. To estimate the *k*-mer combination space, *i.e.* the *k*-mer size, required, we first calculate the probability of *k*-mers with *x* error bases at the front *k-*1 bases as follows:

$p=C_{k-1}^{\chi}E^{\chi}\left( 1-E \right)^{\left( k-x-1 \right)}$	(3)

Where $p$ stands for the probability of a (*k-1)*-mer with$x$ error bases when the error rate is $E$. It should be clarified that *E* refers to the error rate after the exclusion of noise *k*-mers. Generally, the error rates can be decreased for more than one order of magnitude after the exclusion of low coverage *k*-mers. Thus, the error rate after error exclusion should be lower than 0.01 if the strand error rate ≤ 0.1. With *E* = 0.01, the probabilities of a *k*-mer (12 ≤ *k* ≤ 40) containing various error bases at the front *k*-1 bases were obtained based on formula (3). As shown in **Fig. S11**, the probabilities of a *k*-mer (12 ≤ *k* ≤ 40) with three error bases at the front *k*-1 bases is lower than 0.01. This means if we enlarge the *k*-mer size obtained by formula (2) by 3×2 bases, the probability of a noise *k*-mer crash with another noise *k*-mer is lower than 0.01. Thus, to avoid path loops/forks, we enlarge the *k*-mer size estimated by formula 2) by 6, resulting in the following formula for choice of proper *k*-mer size with specific data scale *D* (bits):

$k=\frac{\ln D- \ln2}{\ln4} +7$	(4)

Base on formula (4), the decoding capacity is estimated to be around 1TB with a *k*-mer size of 27. The choices of *k*-mer sizes with data volumes ranging from 1KB to 1EB are listed in **Table S1**.

**Investigation of the potentials of DBGPS by simulation**

The robustness of strand paths in DBG with multiple error-rich sequence copies is analyzed using the script of “*DBGPS_Potentials.py*”. The was performed with a *k*-mer size of ten and a fountain seed of one. For specific rates and types of errors tested, the simulation is iterated 1,000 times. The path conserved rates, which indicate the theoretical maximal strand decoding rates (*S_m_*), are estimated using sequence copies in a range of 3 to 25; Detailed simulation results are provided in **Data S1**.

**Error handling test of DBGPS in comparison with multiple-alignment based strategy**

The performance test results shown in **Fig. 2A-G** were obtained with Python version DBGPS (“*deBruijnGraph.py*”). For the simulations in **Fig. 2A-E**, a strand copy number of 20 and a *k*-mer size of 18 is applied. Results presented in **Fig. 2C-E** were obtained with script “*performance_errors_decoding_rate.py*”. Results presented in **Fig. 2A-B** were obtained with script “*performance_brk_lig_decoding_rate.py*”. For the introduction rearrangements in **Fig. 2B**, rounds of random breaking and ligating were performed to introduce specific rates of rearrangements. For each round, 1% DNA breaks were introduced followed by random pairwise ligation of the obtained DNA fragments. For example, to introduce 3% rearrangements, three rounds of breaking and ligating operations were performed, each round introducing 1% rearrangements. For the introduction mixed errors in **Fig. 2C**, with specific error rate *E*, the strands were introduced with *E*/*3* substitutions, *E*/*6* insertions, *E*/*6* deletions, and *E*/*6* DNA breaks followed by random pairwise ligation of the obtained DNA fragments. The script “*performance_one_many_copies.py*” was utilized for the simulation of the results presented in **Fig. 2F**. A strand error rate of 3% (1.5% substitutions, 0.75% insertions, and 0.75 deletions) and a *k*-mer size of 12 is applied in the simulations of **Fig. 2G**. Muscle 3.8.31^1^ was used for multiple sequence alignments. For exclusion of noise *k*-mers with various sequence copies of $n$, we used the following empirical formula:

$$cutoff=\left\{ \begin{aligned} 0 n<3\text{ } \\ 1 &3\leq n \leq25 \\ \log_{2} n/25+2 n>25 \end{aligned} \right.$$

A Python pipeline was developed for CL-MA based strand reconstruction (<https://github.com/Scilence2022/DBGPS_Python/blob/main/CL-MA-Decoding-test.py>). For each step of CL-MA, the fastest program(s) to the best of our knowledge were employed. For the clustering step, stracode^2^ was applied. For paired-end read assembly, we choose Flash^3^ after testing Pear^4^ and Flash^3^. Our preliminary tests show that Muscle^1^ is faster when the sequence number is below 28, but became slower than Mafft^5^ when introducing more sequences. Therefore, both Muscle and Mafft are utilized in the multiple alignment step to achieve the best performance. For the consensus calling, the majority voting function in Python, utilized in a previous DNA data storage study, was employed ^6^.

**Large-scale simulations**

Large-scale simulations were performed on a server installed with two Intel Xeon (Cascade Lake) Platinum 8269/8269CY CPUs and 1.5 TB of memory. To assess the decoding complexity of DBGPS with large-scale data sets, simulations with input data volumes ranging from 1 MB to 1 GB were performed. The Uniref90 protein database was downloaded and used as input for simulation. For different input scales, we extracted various numbers of protein sequences and used pigz to compress them into a proper size. The DNA fountain codes were employed to generate the DNA strands with a length of 200 bp. Each DNA strand was set to carry 35 bytes of data, 4 bytes of index, 2 bytes of CRC codes and two primer landing sites of 18nt. For each data volume, three independent simulations were performed with degree seeds of 1, 2 and 3. Different numbers of DNA droplets are generated for different data scales with a strand redundancy rate of 50%. The generated DNA droplets were filtered with DBGPS-ft to filter out the entangled strands with a *k*-mer size of 25 and maximal coverage of 1. For input files in sizes of 1 MB to 1 GB, we abstracted filtered droplets for numbers of 3E4 to 3E7. Error-rich strand copies are then simulated with 1.5% substitutions, 0.75% insertions and 0.75% deletions, and strand copy number of 25. Counting of the *k*-mers with large file requires huge memory. The *k*-mer counting step was performed with JellyFish 2.3 ^7^ with a bloom filter which can reduce memory consumption. A *k*-mer size of 27 and a thread number of 100 were applied for *k*-mer counting. The *k*-mers with a coverage >=3 were dumped and then handled by DBGPS for strand reconstruction. To perform *k*-mer counting with JellyFish using a bloom filter, we used a command similar as follows: *jellyfish count [sequence file] -m 27 -t 100 -s 1G --bf-size 64G -o CountResults*. For the data scale of 1 GB, we used a bloom filter size of 64 GB. For other data scales, the bloom filter data size was changed adaptively. To dump the *k*-mers with a coverage ≥3, we used the following command: *jellyfish dump -L 3 CountResults -o CountResultsDumpL3*. To read the *k*-mers and perform DBGPS strand reconstruction, we used a command similar as follow: *DBGPS -k 27 -c 1 -a 100000001 -b 405045800 CountResultsDumpL3 > CountResultsDumpL3.dec*. The decoded strands then can be found in the generated file of “*CountResultsDumpL3.dec*”.

**Error analysis of sequencing data**

The error rates of the sequencing rates are complicated to be accounted, especially for the sequencing results with massive strand breaks and rearrangements. Furthermore, the error rates cannot reflect the data quality straightforwardly. For example, a dataset of sequencing reads may contain higher rates of base errors, but also a small rate of missed strands. Such a dataset should be considered as a high-quality readout although the base error rate is high. For better estimation of the sequencing qualities of DNA data storage and simplification of the error analysis process in the concept of de Bruijn graph theory, here we introduce two indicators: $K_{d}$ (in range of 0 to 1) and $K_{n}$ (in range of 0 to ∞), which together can well define the data quality and are easy to be estimated. The $K_{d}$ stands for the dropout rate of all correct *k*-mers and $K_{n}$ is the ratio of noise *k*-mers and correct *k*-mers. DNA data with a $K_{d}$ and a $K_{n}$ close to 0 stands for high-quality data. Other than error rate which is complicated for accounting, $K_{d}$ and $K_{n}$ can be easily calculated by the following formulas:

$K_{d}=K_{lost}/K_{ori}$	(5)

$K_{n}=K_{all}/K_{corr}$	(6)

In formula (5), $K_{ori}$ stands for the number of *k*-mers in the original strand sequences and $K_{lost}$ stands for the number of *k*-mers that are presented in the original encoded sequences but not presented in the sequencing results. In formula (6), $K_{all}$ stands for the number of *k*-mers in the sequencing results and $K_{corr}$stands for the number of correct *k*-mers, *i.e.* *k*-mers that are presented in the original encoded sequences. DNA data with a $K_{d}$ and a $K_{n}$ close to 0 stands for high-quality data with high data integrity and low noises. The maximal strand decoding rate $S_{m}$ is determined by $K_{d}$. With specific $K_{d}$, the actual strand decoding rate $S_{r}$ is affected by the value of $K_{n}$. The greedy path search becomes extremely slow when decoding DNA data with higher $K_{n}$ values, due to massive branch paths introduced by the noise *k*-mers.

To assess the error rates of sequencing results, we first run simulations to generate strand sequences with specific error rates in ranges of 0.1% to 10% with a step size of 0.1%. With ten strand copies, the $K_{n}$ values of simulated error-rich strand sequences were then calculated respectively. We marked these values as $K_{n10}$ values, where the number 10 stands for an average strand coverage of 10. We then perform polynomial fitting to describe the relationships between $K_{n10}$ values and error rates (**Fig. S12**). With the fitting formula, the error rates of specific retrieval sequencing results then can be easily estimated with the $K_{n10}$ values calculated by random samples of the sequencing reads.

The distribution of the fragment length of the accelerated aging samples was performed by FastP v0.23.2 ^8^ by the function of “insert size” analysis.

The analyze the rearrangements of the three representative samples, the sequencing reads were aligned to the designed strand sequences using BLAST+ 2.2.27^9^ with expected value of 1E5, identity of 98%, and align length of 34 bp. The reads with multiple hits with different strand sequences designed were marked as “reads with rearrangements”.

**Test of block checking codes**

To test the effectiveness of block checking codes, *i.e.*, inserting one parity base every few bases, the sequencing data of file1 from the study by Antkowiak *et al. ^6^* was utilized. The designed sequences were used to simulate the function of the error checking bases. During the simulated “path searching”, the noise *k*-mers with single base error were discarded directly. For the noise *k*-mers with more than one base error were discarded randomly with a probability of ¾. A specialized script “*test_block_codes.py*” to repeat this simulation process was provided at: https://github.com/Scilence2022/DBGPS_Python/.

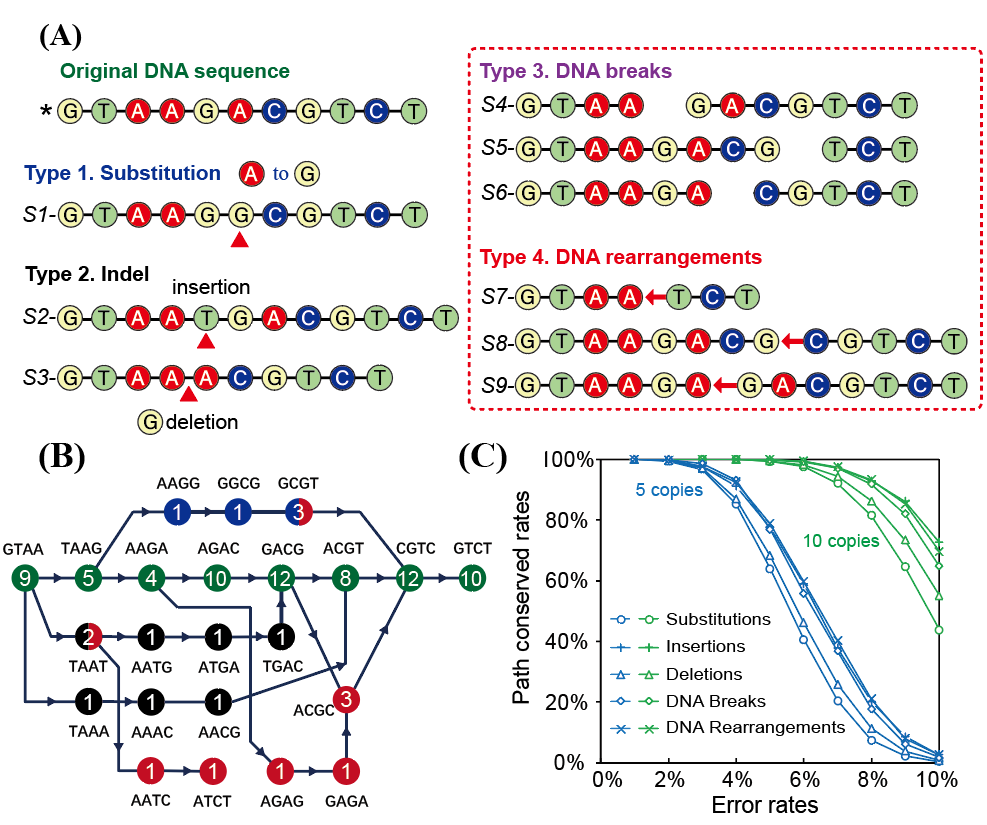

**Fig. S1. Four types of errors in DNA data inner sub-channel and theoretical potentials of de Bruijn Graph-based strand reconstruction.** (A) Illustration of substitutions, indels, DNA breaks and rearrangements in inner sub-channel of DNA data storage. (B) Illustration of de Bruijn graph (DBG) and appearances of the four types of errors in DBG. In graph theory, de Bruijn graph is obtained by taking all strings over any finite alphabet of length *k* as vertices, and adding edges between vertices that have an overlap of *k*−1. The representative DBG was constructed from the nine error-containing sequence copies shown in **Fig. S1A** with a *k*-mer size of four. The numbers inside the circles are frequencies of occurrences, *i.e.* the coverages, of corresponding *k*-mers. (C) Robustness of strand path in DBG with multiple sequence copies containing various types and rates of errors. For specific rates and types of errors tested, the simulation was iterated 1,000 times. The path conserved rates, which indicate the theoretical maximal strand decoding rates (*S_m_*), are estimated using sequence copies of five and ten respectively. Detailed simulation results with sequence copies in a range of 3 to 25 are provided in **Data S1**.

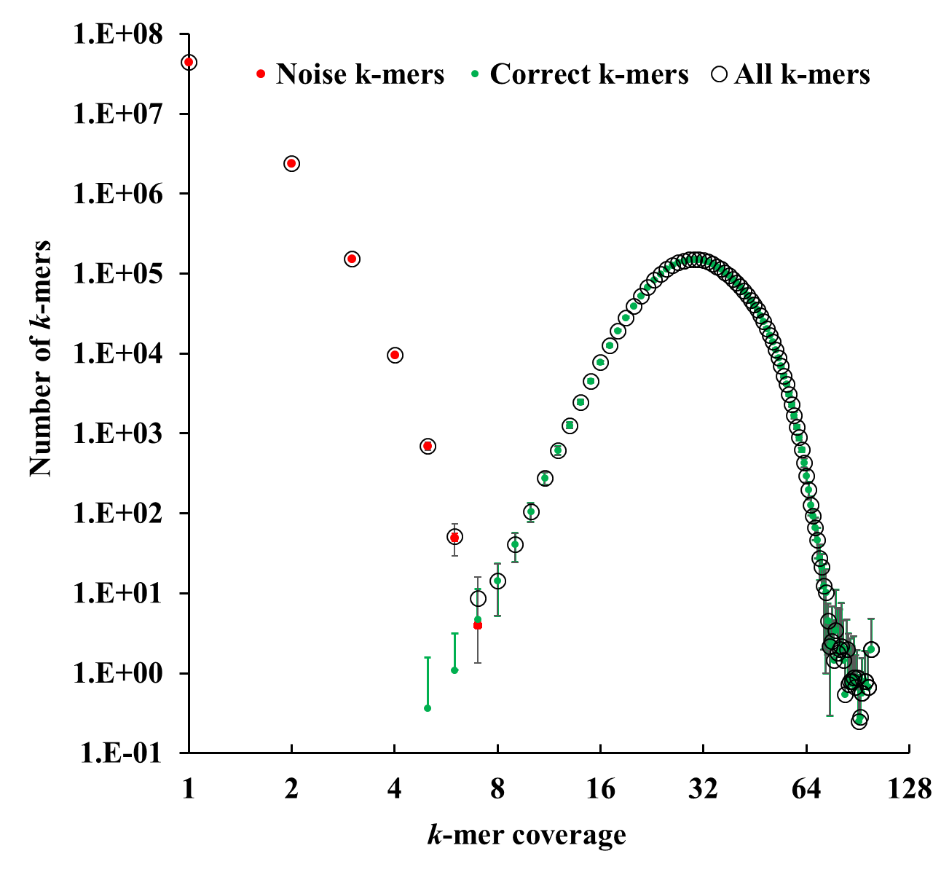

Fig. S2.

**Coverage is a suitable indicator to distinguish noise and correct *k*-mers.**

The simulation was performed with 10,000 DNA droplets and an expected strand copy number of 30. Three individual simulations were performed and the standard deviations were shown.

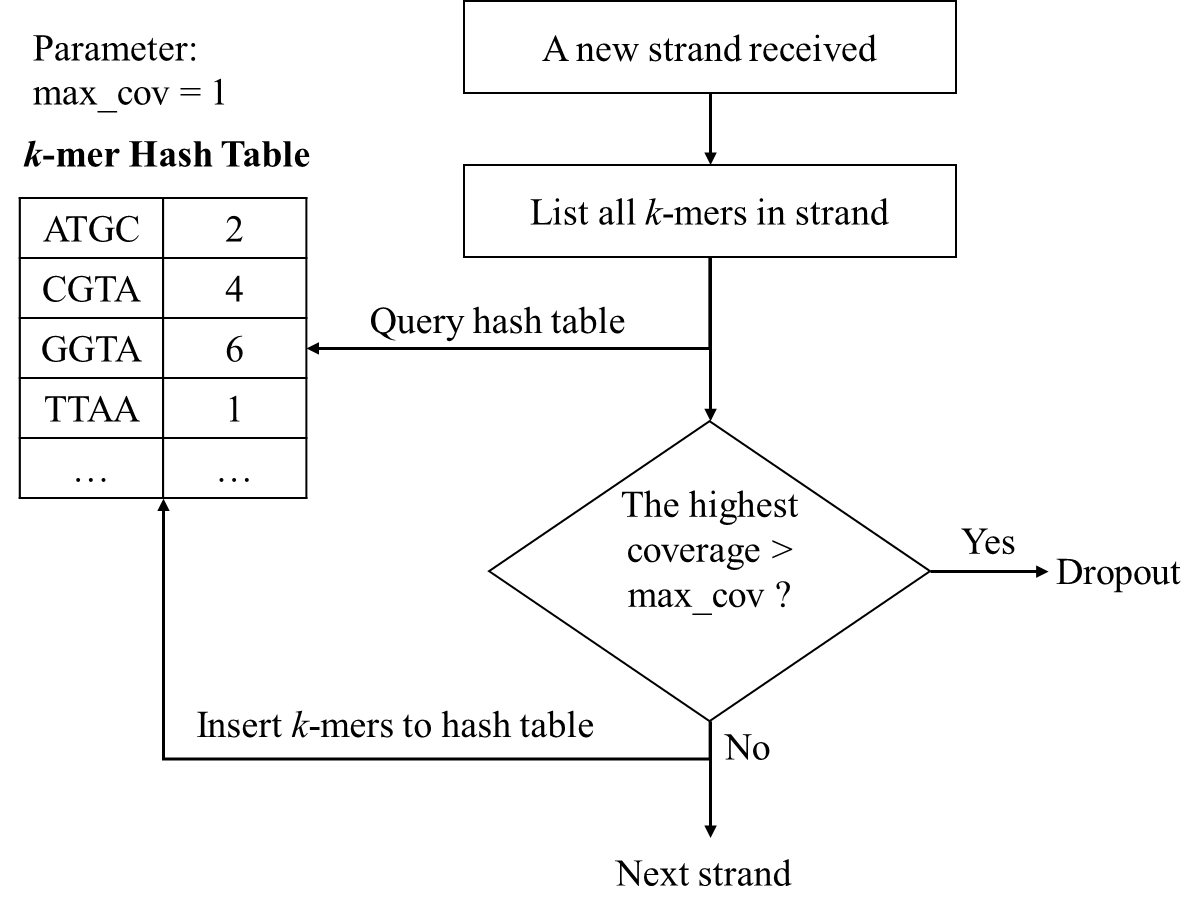

Fig. S3.

**Strand filtering process to avoid strand entanglements in DBG.** With max_cov = 1, all the strands show more than one cross-links with other strands will be filtered.

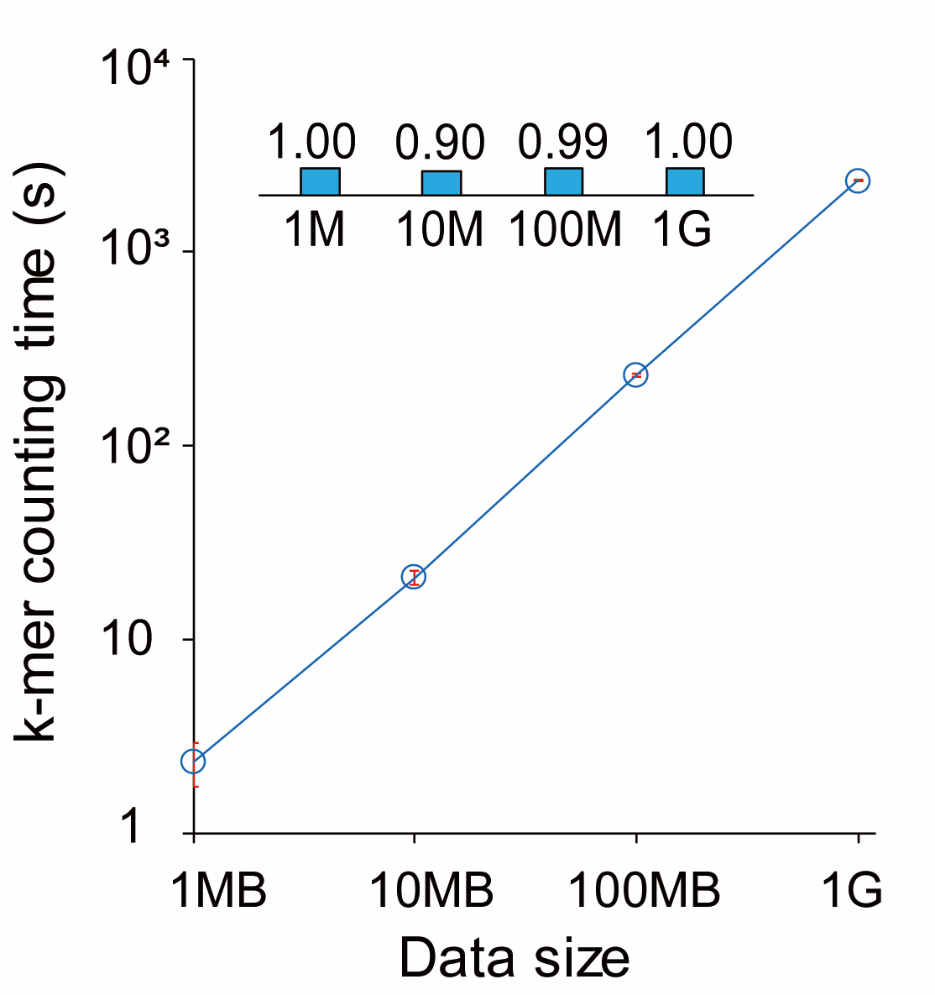

Fig. S4.

***k*-mer counting time with data volumes ranging from 1 MB to 1 GB**. The small bar chart at the top shows the folds of counting time per strand compared to 1 MB scale.

| Data size (MB) | Decoding time (s) |
| --- | --- |
| 1 | 29.8717 |
| 2 | 31.0439 |
| 3 | 32.6586 |
| 4 | 34.5606 |
| 5 | 36.6762 |
| 6 | 38.9621 |
| 7 | 41.3896 |
| 8 | 43.9383 |
| 9 | 46.593 |
| 10 | 49.3419 |
| 11 | 52.1754 |
| 12 | 55.0859 |
| 13 | 58.067 |
| 14 | 61.1131 |
| 15 | 64.2197 |
| 16 | 67.3828 |
| 17 | 70.5987 |
| 18 | 73.8644 |
| 19 | 77.177 |
| 20 | 80.5342 |
| 21 | 83.9337 |
| 22 | 87.3735 |
| 23 | 90.8517 |
| 24 | 94.3668 |
| 25 | 97.917 |
| 26 | 101.501 |
| 27 | 105.118 |
| 28 | 108.766 |
| 29 | 112.444 |
| 30 | 116.151 |
| 31 | 119.887 |
| 32 | 123.649 |
| 33 | 127.439 |
| 34 | 131.253 |
| 35 | 135.093 |
| 36 | 138.957 |
| 37 | 142.844 |
| 38 | 146.755 |
| 39 | 150.687 |
| 40 | 154.641 |
| 41 | 158.616 |
| 42 | 162.612 |
| 43 | 166.628 |
| 44 | 170.664 |
| 45 | 174.719 |
| 46 | 178.793 |
| 47 | 182.885 |
| 48 | 186.995 |
| 49 | 191.123 |
| 50 | 195.268 |
| 51 | 199.43 |
| 52 | 203.608 |
| 53 | 207.803 |
| 54 | 212.014 |
| 55 | 216.24 |
| 56 | 220.482 |
| 57 | 224.739 |
| 58 | 229.01 |
| 59 | 233.297 |
| 60 | 237.597 |
| 61 | 241.912 |
| 62 | 246.24 |
| 63 | 250.582 |
| 64 | 254.938 |
| 65 | 259.307 |
| 66 | 263.689 |
| 67 | 268.083 |
| 68 | 272.491 |
| 69 | 276.91 |
| 70 | 281.342 |
| 71 | 285.786 |
| 72 | 290.242 |
| 73 | 294.71 |
| 74 | 299.189 |
| 75 | 303.68 |
| 76 | 308.182 |
| 77 | 312.695 |
| 78 | 317.219 |
| 79 | 321.754 |
| 80 | 326.3 |
| 81 | 330.856 |
| 82 | 335.422 |
| 83 | 339.999 |
| 84 | 344.586 |
| 85 | 349.184 |
| 86 | 353.791 |
| 87 | 358.408 |
| 88 | 363.034 |
| 89 | 367.671 |
| 90 | 372.317 |
| 91 | 376.972 |
| 92 | 381.636 |
| 93 | 386.31 |
| 94 | 390.993 |
| 95 | 395.684 |
| 96 | 400.385 |
| 97 | 405.095 |
| 98 | 409.813 |
| 99 | 414.54 |
| 100 | 419.275 |
| 101 | 424.019 |
| 102 | 428.771 |
| 103 | 433.532 |
| 104 | 438.3 |
| 105 | 443.077 |
| 106 | 447.862 |
| 107 | 452.655 |
| 108 | 457.456 |
| 109 | 462.264 |
| 110 | 467.08 |
| 111 | 471.905 |
| 112 | 476.736 |
| 113 | 481.575 |
| 114 | 486.422 |
| 115 | 491.276 |
| 116 | 496.138 |
| 117 | 501.006 |
| 118 | 505.882 |
| 119 | 510.765 |
| 120 | 515.656 |
| 121 | 520.553 |
| 122 | 525.457 |
| 123 | 530.369 |
| 124 | 535.287 |
| 125 | 540.212 |
| 126 | 545.143 |
| 127 | 550.082 |
| 128 | 555.027 |
| 129 | 559.978 |
| 130 | 564.936 |
| 131 | 569.901 |
| 132 | 574.872 |
| 133 | 579.85 |
| 134 | 584.834 |
| 135 | 589.824 |
| 136 | 594.821 |
| 137 | 599.823 |
| 138 | 604.832 |
| 139 | 609.847 |
| 140 | 614.868 |
| 141 | 619.895 |
| 142 | 624.929 |
| 143 | 629.968 |
| 144 | 635.013 |
| 145 | 640.063 |
| 146 | 645.12 |
| 147 | 650.183 |
| 148 | 655.251 |
| 149 | 660.325 |
| 150 | 665.405 |
| 151 | 670.49 |
| 152 | 675.581 |
| 153 | 680.677 |
| 154 | 685.779 |
| 155 | 690.887 |
| 156 | 695.999 |
| 157 | 701.118 |
| 158 | 706.242 |
| 159 | 711.371 |
| 160 | 716.505 |
| 161 | 721.645 |
| 162 | 726.79 |
| 163 | 731.94 |
| 164 | 737.095 |
| 165 | 742.256 |
| 166 | 747.421 |
| 167 | 752.592 |
| 168 | 757.768 |
| 169 | 762.948 |
| 170 | 768.134 |
| 171 | 773.325 |
| 172 | 778.521 |
| 173 | 783.721 |
| 174 | 788.927 |
| 175 | 794.137 |
| 176 | 799.353 |
| 177 | 804.573 |
| 178 | 809.797 |
| 179 | 815.027 |
| 180 | 820.261 |
| 181 | 825.5 |
| 182 | 830.744 |
| 183 | 835.992 |
| 184 | 841.245 |
| 185 | 846.503 |
| 186 | 851.765 |
| 187 | 857.031 |
| 188 | 862.302 |
| 189 | 867.578 |
| 190 | 872.858 |
| 191 | 878.143 |
| 192 | 883.432 |
| 193 | 888.725 |
| 194 | 894.023 |
| 195 | 899.325 |
| 196 | 904.632 |
| 197 | 909.942 |
| 198 | 915.257 |
| 199 | 920.577 |
| 200 | 925.9 |

| Data size (MB) | Decoding time (s) |
| --- | --- |
| 1 | 2.36 |
| 10 | 31.27 |
| 100 | 468.05 |
| 1000 | 5867.73 |

Fig. S5.

**Decoding complexity of DBGPS estimated by curve fitting experiments with large-scale simulation results.** The decoding time curve of DBGPS shows high consistency to a complexity of $O(nlogn)$, where *n* stands for the data size.

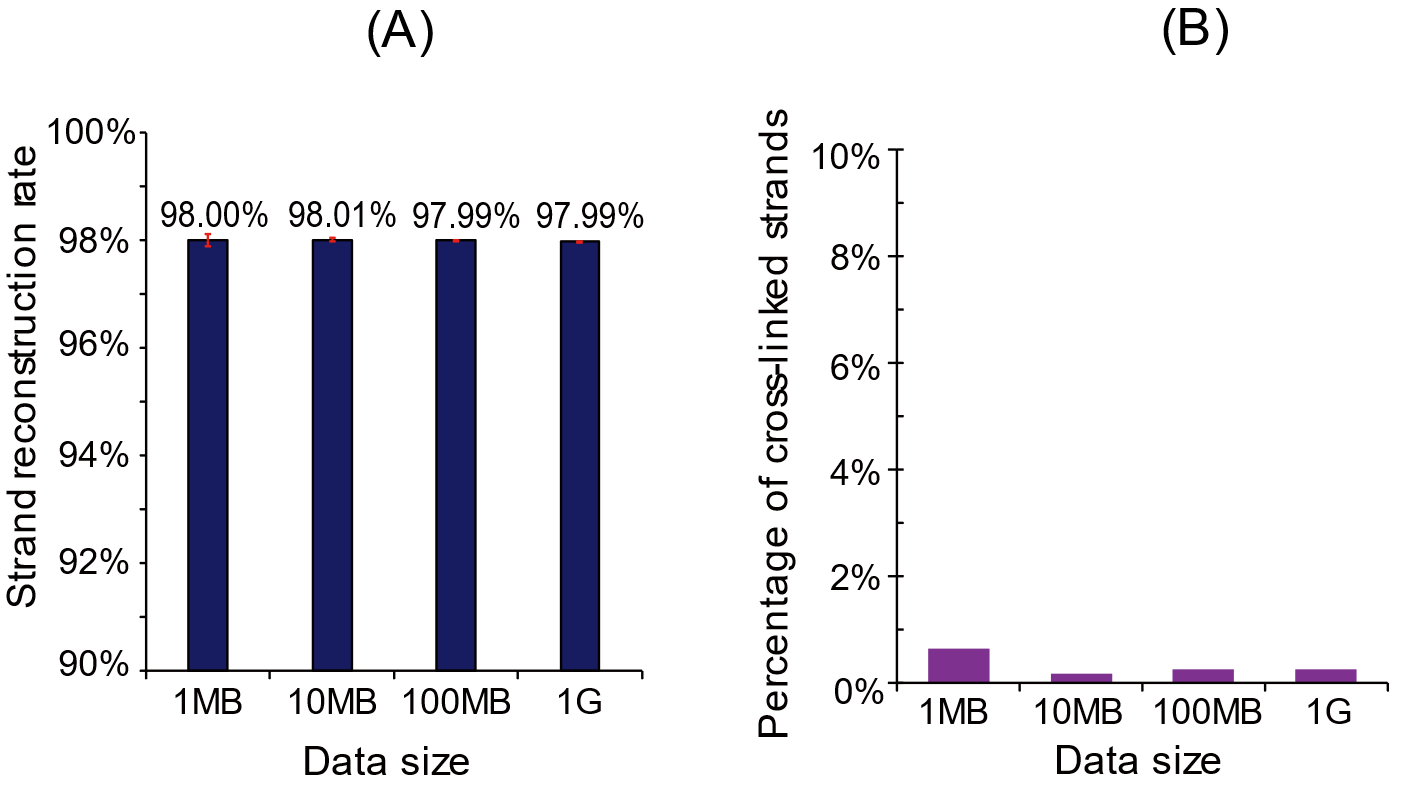

Fig. S6.

**Large-scale simulation analysis.** (A). Strand reconstruction rates by DBGPS with data volumes ranging from 1 MB to 1 GB. (K). Cross-link analysis of the DBG derived from input data with volumes ranging from 1 MB to 1 GB. The y-axis displays the percentages of strands with cross-links to other strands in the corresponding DBG. The strand cross-links are formed by the repeated presentation of (*k*-1)-mer fragments in different strands.

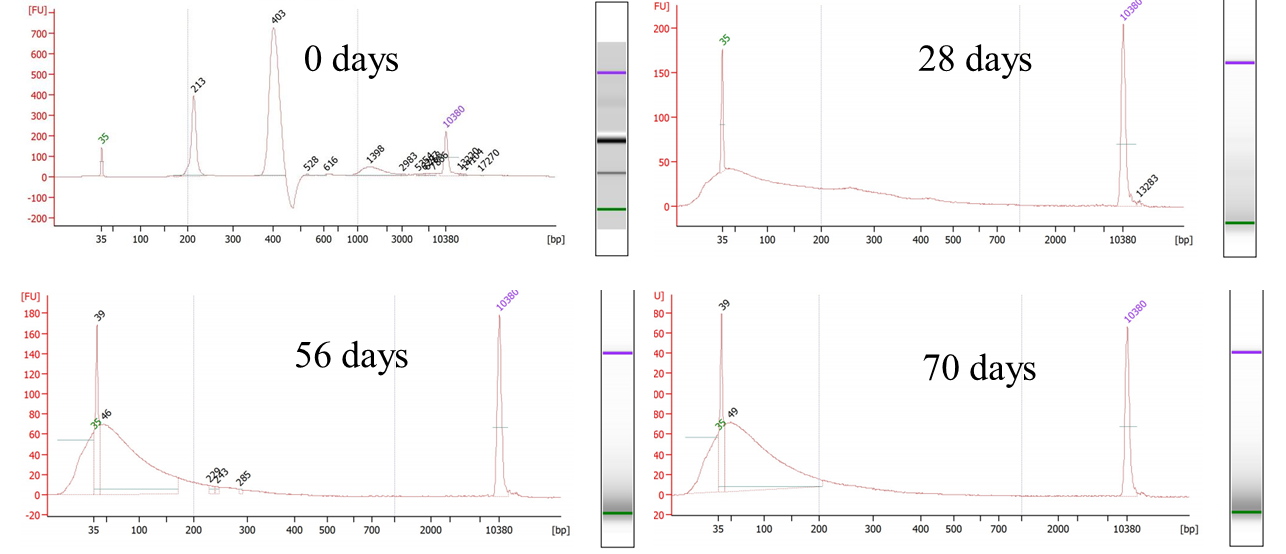

Fig. S7.

**Agilent 2100 Bioanalyzer analysis of the accelerated aged samples which have been treated at 70°C for 0 to 70 days.** The peaks of 35 and 10380 are formed by the internal standards.

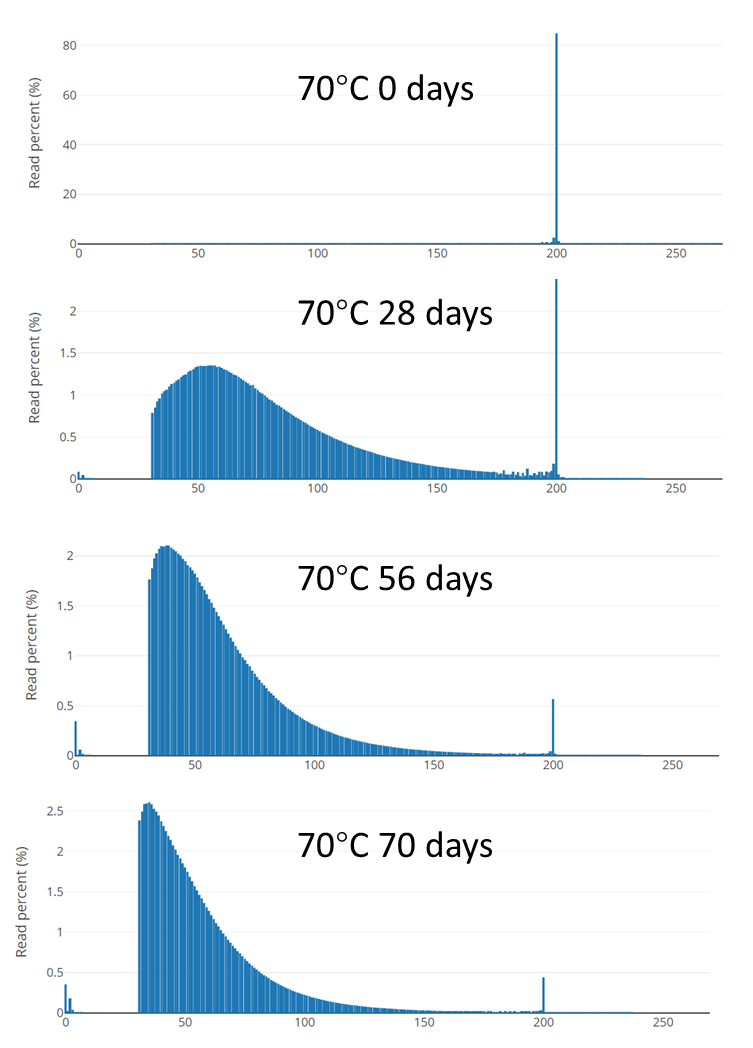

Fig. S8.

**Insert size distribution of the accelerated aged samples based on sequencing results.**

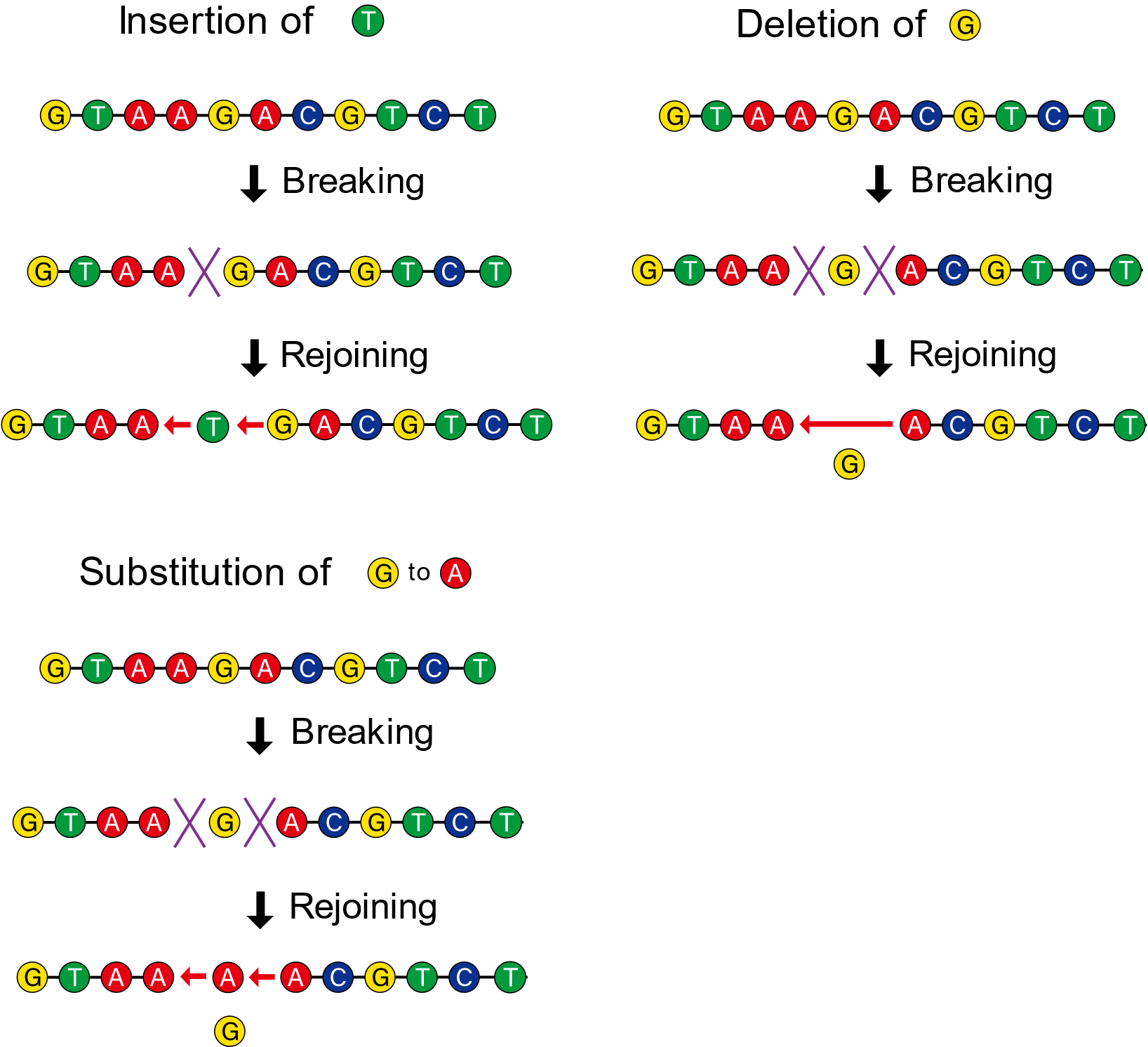

Fig. S9.

**Illustration of indel and substitution as special cases of “DNA rearrangements”.**

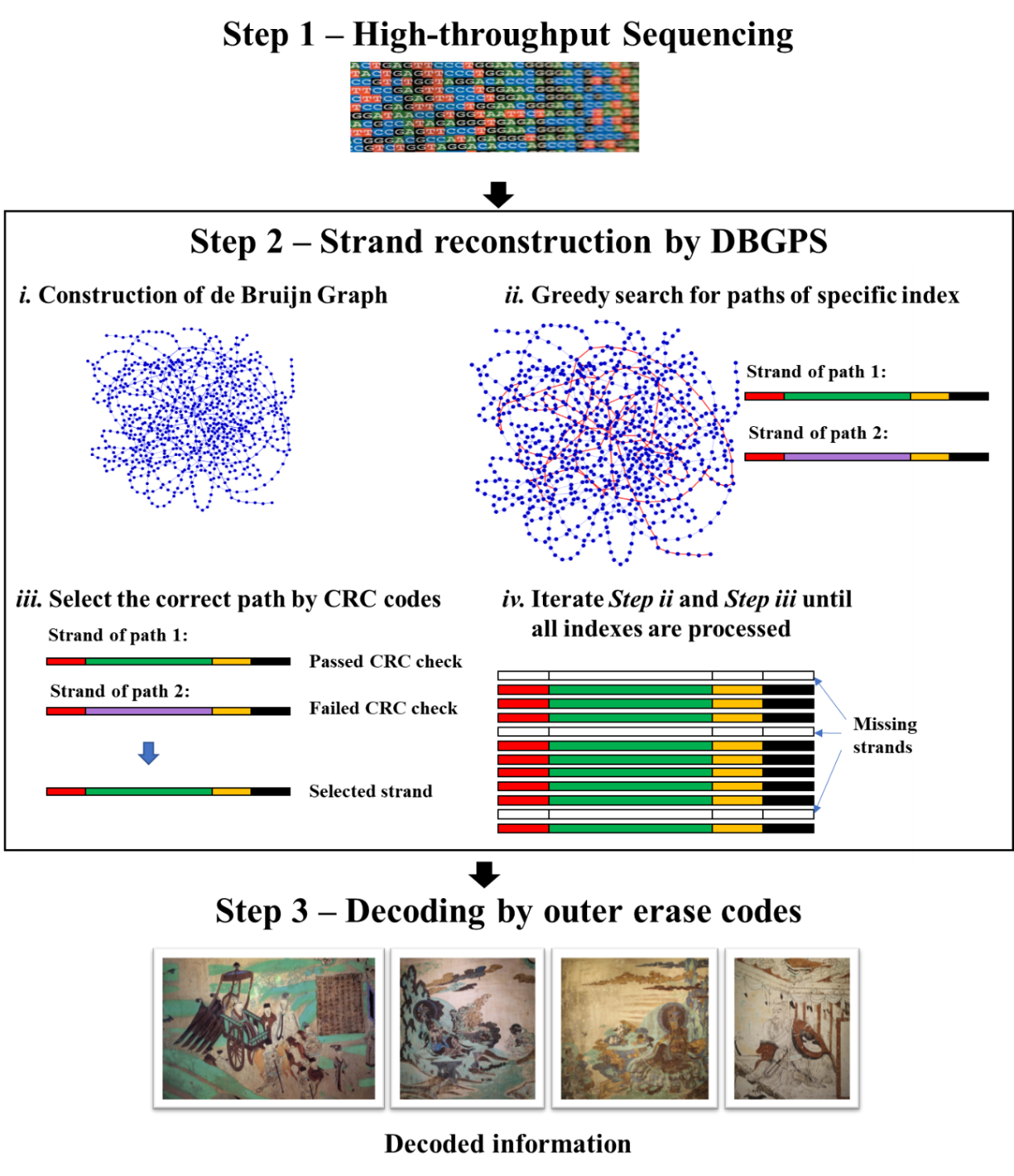

Fig. S10.

**Integration of DBGPS with outer erasure codes for accurate data decoding.**

Probability of k-mers with x error bases

| Number of error bases (x) | k=40 | k=36 | k=34 | k=30 | k=24 | k=20 | k=16 | k=12 |
| --- | --- | --- | --- | --- | --- | --- | --- | --- |
| 1 | 0.266196 | 0.248694 | 0.229307 | 0.207912 | 0.184375 | 0.158558 | 0.130312 | 0.099482 |
| 2 | 0.0510882 | 0.042705 | 0.0347435 | 0.0273015 | 0.0204861 | 0.0144143 | 0.00921397 | 0.00502434 |
| 3 | 0.00636452 | 0.004745 | 0.00339246 | 0.00229811 | 0.00144851 | 0.000825063 | 0.000403305 | 0.000152253 |
| 4 | 0.000578593 | 0.000383434 | 0.000239871 | 0.000139279 | 7.31572e-05 | 3.33359e-05 | 1.22214e-05 | 3.07582e-06 |
| 5 | 4.09106e-05 | 2.4013e-05 | 1.30839e-05 | 6.47156e-06 | 2.80806e-06 | 1.01018e-06 | 2.71586e-07 | 4.34964e-08 |
| 6 | 2.34168e-06 | 1.21278e-06 | 5.72695e-07 | 2.39688e-07 | 8.50926e-08 | 2.38089e-08 | 4.57215e-09 | 4.39357e-10 |
| 7 | 1.11509e-07 | 5.07512e-08 | 2.066e-08 | 7.26326e-09 | 2.08741e-09 | 4.46632e-10 | 5.93786e-11 | 3.16997e-12 |
| 8 | 4.5054e-09 | 1.79424e-09 | 6.2606e-10 | 1.83416e-10 | 4.21699e-11 | 6.76715e-12 | 5.99784e-13 | 1.60099e-14 |
| 9 | 1.56754e-10 | 5.43708e-11 | 1.61609e-11 | 3.91122e-12 | 7.0993e-13 | 8.3545e-14 | 4.71211e-15 | 5.39055e-17 |

Fig. S11.

**Probabilities of specific size *k*-mers containing x error bases at the front *k*-1 bases.**

This figure was drawn based on the formula of $p=C_{k-1}^{\chi}E^{\chi}\left( 1-E \right)^{\left( k-x-1 \right)}$ with an *E* value of 0.01.

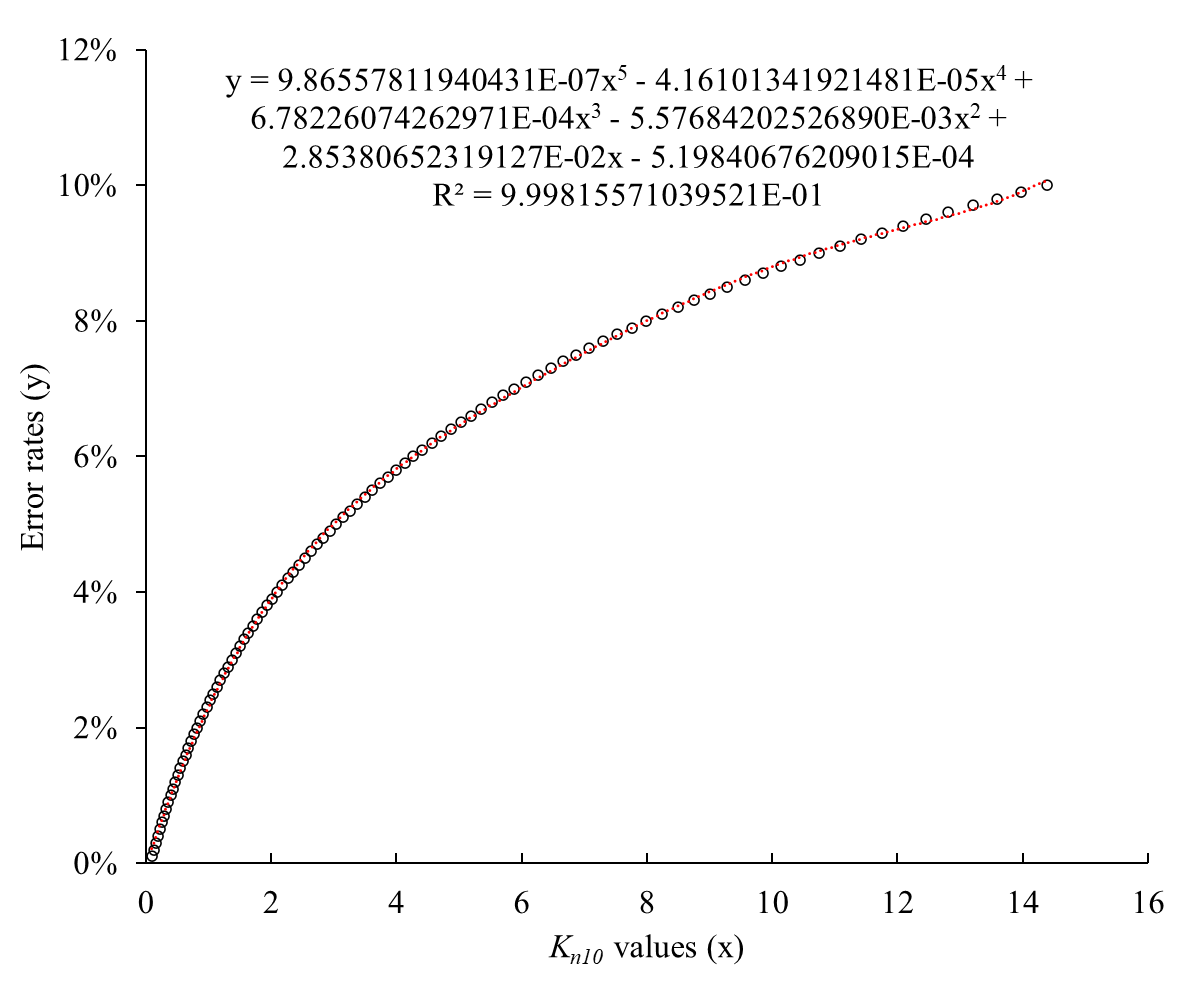

Fig. S12.

**Polynomial fitting results of *K_n10_* values and error rates.**

The obtained formula was used for estimation of the error rates using *K_n10_* values obtained by random sampling of the sequencing reads.

| Data scale | 1KB | 1MB | 1GB | 1TB | 1EB |
| --- | --- | --- | --- | --- | --- |
| Choice of *k*-mer size | ≥12 | ≥17 | ≥23 | ≥27 | ≥32 |

Table S1.

**Choice of *k*-mer sizes with various data volumes ranging from 1KB to 1EB estimated with an error rate of 0.01.** *Note: Here, the error rate refers to the error rate after exclusion of noise k-mers.*

|  | Number of reconstructed strands | *S_r_* |
| --- | --- | --- |
| 70°C-28 days | 166,784 | 79.42% |
| 70°C-56 days | 86,939 | 41.40% |
| 70°C-70 days | 38,189 | 18.19% |
| Multiple Retrieval-Sample A | 24,660 | 11.74% |
| Multiple Retrieval-Sample B | 18,092 | 8.62% |
| Multiple Retrieval-Sample C | 19,674 | 9.37% |
| ePCR#1 | 208,488 | 99.28% |
| ePCR#2 | 207,438 | 98.78% |
| ePCR#3 | 164,459 | 78.31% |
| ePCR#4 | 55,703 | 26.53% |
| ePCR#5 | 35,208 | 16.77% |
| ePCR#6 | 1,158 | 0.55% |

**Table S2. Strand reconstruction results with the sequencing data of the three harsh experiments by CL-MA method.**

|  | Estimated error rate | Average length | Percentages of corrupted strands | *S_m_* | Number of reconstructed strands | *S_r_* |
| --- | --- | --- | --- | --- | --- | --- |
| 70°C-0 days | 0.18% | 196.25 bp | 0.25% | 99.75% | 208,833 | 99.44% |
| 70°C-28 days | 1.82% | 80.91 bp | 13.81% | 99.36% | 207,642 | 98.88% |
| 70°C-56 days | 2.66% | 62.28 bp | 54.58% | 99.05% | 205,883 | 98.04% |
| 70°C-70 days | 2.69% | 57.86 bp | 80.82% | 98.35% | 202,469 | 96.41% |

Table S3.

**Strand reconstruction details of the accelerated aging samples by DBGPS.**

|  | Estimated error rate | *S_m_* | Number of reconstructed strands | *S_r_* |
| --- | --- | --- | --- | --- |
| ePCR#1 | 0.53% | 99.85% | 208,937 | 99.49% |
| ePCR#2 | 0.62% | 99.65% | 208,014 | 99.05% |
| ePCR#3 | 1.22% | 99.20% | 205,310 | 97.77% |
| ePCR#4 | 1.86% | 99.18% | 202,831 | 96.59% |
| ePCR#5 | 3.44% | 98.22% | 192,916 | 91.86% |
| ePCR#6 | 6.05% | 94.67% | 160,791 | 76.57% |

Table S4.

**Strand reconstruction details of the error-prone PCR samples by DBGPS.**

|  |  | *S_m_* | Number of reconstructed strands | *S_r_* |
| --- | --- | --- | --- | --- |
| Control |  | 98.55% | 206,234 | 98.21% |

Table S5.

**Data retrieval details of a diluted sample with a physical density of 295 PB/g (~1000 molecular copies).**

| Block length (bp) | Number of reconstructed strands (Simulated) | *S_r_*  (Simulated) |
| --- | --- | --- |
| 3 | 16,311 | 99.6% |
| 4 | 16,306 | 99.5% |
| 5 | 16,275 | 99.3% |
| 6 | 16,239 | 99.1% |
| 7 | 16,202 | 98.9% |
| 8 | 16,184 | 98.8% |
| 9 | 16,140 | 98.5% |
| 10 | 16,104 | 98.3% |

Table S6.

**Performance of DBGPS and block parity checking in low cost, chip-based oligo synthesis of low quality.** The results are estimated by a simulated greedy path search process using the sequencing data of file 1 in the study by Antkowiak *et al.* ^6^. A *k*-mer size of 18 was applied during simulation. The *k*-mers with a coverage lower than 2 were excluded during simulation. Since this block parity checking mechanism can accurately recognize the noise *k*-mers with single base error. The *k*-mers with single base error were directly removed during the simulated process of greedy path search. The noise *k*-mers with two and more base errors were removed randomly during path search simulation with a probability of 75%.

Movie S1.

End-to-end presentation of the decoding process by DBGPS and outer fountain codes.

The decoding started with raw sequencing reads encoding a 6.8 MB zipped file of ten pictures of Dunhuang murals. The decoding includes two major steps. At first, the raw sequencing reads are handled by DBGPS, generating strand sequences. Then, the DNA fountain codes are then utilized to decipher the original data. At the end of the movie, the decoded file was checked by comparing the MD5 hash values and opening the unzipped pictures manually.

Data S1. (separate file)

Detailed simulation results about the potentials of de Bruijn graph-based strand reconstruction with multiple error-containing strand copies.

Data S2. (separate file)

A zipped file in a size of 6.8 MB used as input data in the robustness verification experiments.

Data S3. (separate file)

The design details of the 21,000 DNA strands encoding the 6.8 MB zipped file.

Data S4. (separate file)

Strand reconstruction details of the 100 independent data retrievals.

**References**

1.	Edgar, R. C. MUSCLE: multiple sequence alignment with high accuracy and high throughput. *Nucleic acids research* **32,** 1792–1797; 10.1093/nar/gkh340 (2004).

2.	Zorita, E., Cuscó, P. & Filion, G. J. Starcode: sequence clustering based on all-pairs search. *Bioinformatics (Oxford, England)* **31,** 1913–1919; 10.1093/bioinformatics/btv053 (2015).

3.	Magoč, T. & Salzberg, S. L. FLASH: fast length adjustment of short reads to improve genome assemblies. *Bioinformatics (Oxford, England)* **27,** 2957–2963; 10.1093/bioinformatics/btr507 (2011).

4.	Zhang, J., Kobert, K., Flouri, T. & Stamatakis, A. PEAR: a fast and accurate Illumina Paired-End reAd mergeR. *Bioinformatics (Oxford, England)* **30,** 614–620; 10.1093/bioinformatics/btt593 (2014).

5.	Katoh, K., Misawa, K., Kuma, K.-i. & Miyata, T. MAFFT: a novel method for rapid multiple sequence alignment based on fast Fourier transform. *Nucleic acids research* **30,** 3059–3066; 10.1093/nar/gkf436 (2002).

6.	Antkowiak, P. L. *et al.* Low cost DNA data storage using photolithographic synthesis and advanced information reconstruction and error correction. *Nature communications* **11,** 5345; 10.1038/s41467-020-19148-3 (2020).

7.	Marçais, G. & Kingsford, C. A fast, lock-free approach for efficient parallel counting of occurrences of k-mers. *Bioinformatics (Oxford, England)* **27,** 764–770; 10.1093/bioinformatics/btr011 (2011).

8.	Chen, S., Zhou, Y., Chen, Y. & Gu, J. fastp: an ultra-fast all-in-one FASTQ preprocessor. *Bioinformatics (Oxford, England)* **34,** i884-i890; 10.1093/bioinformatics/bty560 (2018).

9.	Camacho, C. *et al.* BLAST+: architecture and applications. *BMC bioinformatics* **10,** 421; 10.1186/1471-2105-10-421 (2009).
